# Transcriptomic landscape of Gallbladder cancer reveals altered pathways related to cell cycle and Aurora kinase

**DOI:** 10.1101/2025.09.04.674273

**Authors:** Sajib Kumar Sarkar, Arnab Nayek, Rashmi Minocha, Gurpreet Singh, Deepak Kumar, Nidhi Bharadwaj, Prasenjit Das, Nihar Ranjan Dash, Kailash Kurdia, Abhibroto Karmakar, Ruby Dhar, Subhradip Karmakar

## Abstract

**Background:** Gallbladder cancer (GBC) is a rare but aggressive biliary tract malignancy. This study explores the transcriptomic profile of GBC to identify differentially expressed genes (DEGs) and dysregulated pathways involved in its pathogenesis.

**Methods:** RNA sequencing was performed on 13 GBC tumors and 6 matched controls. Functional enrichment analysis as well as WGCNA were used to identify dysregulated pathways, functionally relevant gene modules and hub genes. Key targets were validated in patient tissues and cell lines.

**Results:** A total of 621 DEGs were identified (247 upregulated, 374 downregulated). Gene set enrichment analysis revealed activation of E2F targets and G2/M checkpoint, with downregulation of bile acid metabolism and estrogen response pathways. A tumor grade-correlated WGCNA module was enriched in cell cycle genes. TPX2 emerged as a central hub gene. Inhibitors of aurora kinase, TPX2 dependent enzyme, significantly reduced proliferation, migration, and invasion in GBC cells. High-grade tumors confirmed elevated Aurora kinase expression.

**Conclusions:** This first transcriptomic analysis of GBC in South-East Asian Indians uncovers key drivers like TPX2 and Aurora kinases in disease progression. The study highlights cell cycle dysregulation and sex-linked signatures, offering insights for biomarker discovery and targeted therapies.

## Background

Gallbladder cancer (GBC) is a highly aggressive biliary tract malignancy with a dismal prognosis. Late-stage detection due to vague clinical presentations and lack of biomarkers results in 5-15% five-year survival rates and a median survival of under 6 months for metastatic disease (1). Only 10% of patients qualify for curative surgery, and standard treatments (surgery, gemcitabine chemotherapy, radiation) show limited efficacy. The cancer’s molecular heterogeneity, chemoresistance, anatomical complexity, and high recurrence rates make it exceptionally challenging to treat (2,3).

Understanding GBC’s molecular pathogenesis through transcriptomic analysis is crucial for identifying altered gene expression profiles, proliferative pathways, and disrupted apoptotic mechanisms. This research could elucidate oncogenic mechanisms and guide the development of novel biomarkers and targeted therapies for this formidable cancer.

High-throughput RNA sequencing (RNA-seq) has transformed cancer research by providing comprehensive transcriptomic profiles that reveal key pathways and hub genes in involved in tumorigenesis and tumor progression. Since chronic cholecystitis is an important risk factor for GBC (4), early identification of at-risk individuals is crucial for manageable intervention.

This study conducted comparative transcriptomic analysis between GBC and chronic cholecystitis cases to understand GBC pathogenesis. We employed weighted gene co-expression network analysis (WGCNA). This systems biology approach transforms complex gene expression data into biologically meaningful network architectures using hierarchical clustering and topological overlap matrices.

WGCNA groups genes into co-expression modules and identifies hub genes that significantly influence cellular signaling pathways. This approach has proven highly effective in cancer research, successfully identifying coregulated gene networks in hepatocellular carcinoma (5) and glioblastoma (6), demonstrating its precision in deciphering cancer-related molecular mechanisms.

WGCNA identifies highly interconnected gene modules corresponding to specific biological processes or cancer hallmarks, assigning an eigengene (first principal component) to each module that correlates with clinical traits to identify disease-relevant networks. Hub genes with high intramodular connectivity represent key regulatory nodes in cancer progression, making this approach valuable for identifying cancer subtypes and predicting patient outcomes (7).

This study utilized RNA-seq and WGCNA to explore GBC’s transcriptomic profile, identifying dysregulated pathways and hub genes involved in disease pathogenesis. Top targets were experimentally validated to confirm their biological significance.

## Methods

### Tissue collection & RNA isolation

Gallbladder control (n=6) and tumor tissues (n=13) were collected from specimens of simple cholecystectomy and extended cholecystectomy. After PBS washing and necrotic tissue removal, 3-5mm pieces were prepared and stored in RNAlater at - 20°C or processed immediately (Supplementary figure S1a).

Tissues were homogenized in Trizol using mortar and pestle. RNA isolation involved adding chloroform to Trizol lysate, phase separation by centrifugation (10,000 RPM, 20 min, 4°C), and column purification of the aqueous phase with isopropanol.

### RNA sequencing

RNA quality was assessed using a Nanodrop spectrophotometer (Thermofisher Scientific, MA, USA), agarose gel electrophoresis and a Bioanalyzer (Agilent Technologies, Inc, CA, USA). Samples with an RNA Integrity Number (RIN) >7 was selected for further RNA-seq analysis (Supplementary figure S1b,c).

The library preparation for RNA-seq was done using Trueseq standard total RNA (Illumina # 20020597) following the manufacturer’s protocol. The final libraries were quantified using Qubit 4.0 fluorometer (Thermofisher #Q33238) using DNA HS assay kit (Thermofisher #Q32851). Sequencing was performed on Illumina NovaSeq 6000 V1.5 platform. Paired-end sequencing (2x150bp) was carried out to achieve >50 million reads from each sample. Read quality was assessed by FastQC. Trimmomatic was used to remove adapters and low-quality bases (<Q30). Filtered reads were aligned to the human reference genome (GRCh38.p13) using STAR, sorted with Samtools, and assembled with StringTie. Gene expression counts were obtained using FeatureCounts. Differential gene expression analysis was employed DESeq2 with transcripts having ≥10 reads in either group. Differentially expressed genes met the criteria of p-value ≤0.05 and |logFC| ≥1 (Supplementary figure S2).

### Functional enrichment analysis of differentially expressed genes (DEGs)

To investigate the biological functions of DEGs, the Gene Ontology (GO) enrichment analysis, Kyoto Encyclopedia of Genes and Genomes (KEGG) pathway analysis, and Gene Set Enrichment Analysis (GSEA) were performed using R packages - “clusterProfiler” (v3.14.3), “enrichplot” (v1.20.0), “ReactomePA”, (v1.44.0), “ggplot2” (v3.4.2) and “msigdbr” (v7.5.1). Molecular Signature Database (MsigDB), GO, and KEGG databases were used for annotation and enrichment for the queried DEGs. A p-value ≤ 0.05 was considered statistically significant.

### Co-expression network construction

Co-expression networks were constructed using WGCNA R package with annotated variable genes. A Pearson correlation matrix was built between all gene pairs, and an appropriate weighting coefficient β was selected for scale-free topology. The correlation matrix was transformed into an adjacency matrix, then a topological overlap matrix (TOM). Hierarchical clustering used average linkage based on TOM dissimilarity, with modules identified by dynamic tree-cutting (minimum size 30). Module eigengenes (first principal components) represented overall module expression levels. High-similarity modules were merged using a threshold of 0.25 based on Pearson correlations between module eigengenes.

### Identification of clinically significant modules

Module-clinical trait correlations were assessed using two methods to identify clinically significant modules. First, Pearson correlation coefficients between clinical traits and module eigengenes (MEs) determined significant associations (p < 0.05). Second, correlations between clinical traits and individual gene expression levels (gene significance, GS) were calculated, with mean absolute GS values representing module-trait association strength. Module Overrepresentation Analysis (ORA) using hypergeometric testing explored the biological significance of identified modules.

### Identification of hub genes

Hub genes were identified as highly interconnected genes with functional importance within modules. Module membership (MM) was calculated by correlating gene expression with module eigengenes (MEs), while intramodular connectivity (IC) represented the sum of correlation coefficients with adjacent genes in the module. Higher IC genes showed greater MM values, indicating crucial roles.

The top 20 genes with highest IC from the most clinically significant module were selected as central genes (MM > 0.6, q-weighted cutoff < 0.001). Gene-gene networks were visualized using Cytoscape, with hub genes defined as those with the highest degree within key networks.

### Functional enrichment analysis of clinically significant modules

To investigate the biological functions and associated signaling pathways of the genes within the most significant clinical module, functional enrichment analysis, including Gene Ontology (GO) enriched terms and Kyoto Encyclopedia of Genes and Genomes (KEGG) pathway analysis, was performed using the clusterProfiler R package (version 4.12.0). Genes from the clinically high significant module with MM > 0.6 and q-weighted < 0.001 were selected for functional enrichment analysis. For the identification of significantly enriched GO terms, KEGG, and ReactomePA pathways, the threshold criteria were set as p < 0.05. Supplementary figure S3 outlines the analysis workflow of WGCNA used for the study.

### Survival Analysis

The prognostic value of hub genes was assessed using TCGA database for cholangiocarcinoma (CHOL). Kaplan-Meier analysis was conducted with the ‘survival’ package in R (version 3.2-3). P values <0.05 were considered significant.

### Immunohistochemistry

Tissue sections were fixed in formalin, deparaffinized with xylene, dehydrated with acetone, and rehydrated with ethanol. Antigen retrieval was performed using heat-induced epitope retrieval (HIER) with citrate or Tris-EDTA buffer at 100°C for 25 minutes. After antibody incubation, DAB substrate developed signals, and hematoxylin counterstaining was performed. GBC (n=10) and control (n=10) sections were analyzed using brightfield microscopy, with image analysis in ImageJ software to compare mean intensities.

### Cell Line Treatment

G415 gallbladder cancer cell-line (Riken Cell Bank, Japan) was used in the study after STR validation (Supplementary figure S4). The cells were treated with 100nM Aurora Kinase A and B inhibitors for 24 hours. Cellular properties and gene expression profiles were assessed post-treatment.

### RNA Isolation and qPCR

RNA was isolated using a column-based method (Promega), and cDNA was synthesized for qPCR with SYBRGreen (Thermofisher). Relative expression was calculated for each target both in cases and controls with respect to GAPDH using the formula

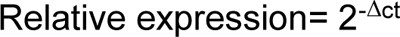

where, ΔCt value of sample = (ct value of target gene - ct value of GAPDH)

Additionally, fold change was calculated for in vitro experiments by Livak’s method where,

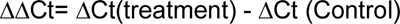

And fold change in expression of a certain gene in the treatment group: 2^(-ΔΔCt)^

Details of all the primers used in the study have been provided in Additional file 2 (Table S1).

### Wound Healing Assay

G415 cells were seeded in 6-well plates (2.5 × 10⁵ cells/well) and scratched once confluent. The wound area was monitored for 24 hours, and closure was analyzed using live cell imaging (Axion BioSystems). Wound closure percentage was calculated as:

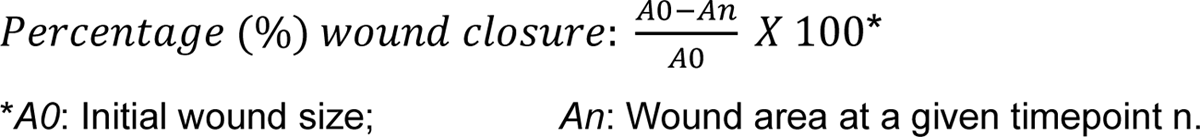

### Cell Cycle Analysis

Cells were fixed and stained with propidium iodide. Data was acquired using a flow cytometer (Beckton Dickinson).

### Statistical Analysis

Statistical analysis was performed using GraphPad Prism (version 9). Quantitative qPCR data were compared using the Mann Whitney U test for two groups or Kruskal-Wallis test with Dunn’s post hoc for multiple comparisons. Data are presented as median ± interquartile range or mean ± standard deviation, with P<0.05 considered significant.

## Results

### Demography and clinical profile of the participants

For RNA sequencing analysis, a total of 19 participants were enrolled in the study, including 13 cases of GBC and 6 with CC. Comprehensive demographic and clinical characteristics of all participants, including age, gender, histopathological findings, and relevant clinical parameters, are summarized in Table 1.

**Table 1:** Demography and clinical profile of GBC cases participated in the study.

| Sl. No. | Characteristics | Number of cases | Percentage |
| --- | --- | --- | --- |
| 1 | **Sex** |  |  |
|  | Female | 7 | 53.8 |
|  | Male | 6 | 46.2 |
| 2 | **Age distribution** |  |  |
|  | Median | 50 |  |
|  | Range | 30-72 |  |
| 3 | **Geographic setting** |  |  |
|  | Rural | 8 | 61.5 |
|  | Urban | 5 | 38.5 |
| 4 | **Localization** |  |  |
|  | Fundus | 7 | 53.8 |
|  | Body | 5 | 38.5 |
|  | Neck | 1 | 7.7 |
| 5 | **Tumor size** |  |  |
|  | T1 | 1 | 7.7 |
|  | T2 | 3 | 23.1 |
|  | T3 | 7 | 53.8 |
|  | T4 | 2 | 15.4 |
| 6 | **Lymph node** |  |  |
|  | N0 | 5 | 38.5 |
|  | N1 | 5 | 38.5 |
|  | N2 | 3 | 23.1 |
| 7 | **Stage** |  |  |
|  | I | 2 | 15.4 |
|  | II | 1 | 7.7 |
|  | III | 6 | 46.2 |
|  | IV | 4 | 30.8 |
| 8 | **Grade** |  |  |
|  | WDADC* | 2 | 15.4 |
|  | MDADC** | 8 | 61.5 |
|  | PDADC*** | 3 | 23.1 |
| 9 | **Tobacco Usage**<br>**(Smoking or chewing)** |  |  |
|  | Yes | 9 | 69.2 |
|  | No | 4 | 30.8 |
| 10 | **Alcohol intake** |  |  |
|  | Yes | 4 | 30.8 |
|  | No | 9 | 69.2 |
| 11 | **Associated gallstone** |  |  |
|  | Yes | 6 | 46.2 |
|  | No | 7 | 53.8 |

\*WDADC- Well differentiated adenocarcinoma
\*\*MDADC- Moderately differentiated adenocarcinoma
\*\*\*PDADC- Poorly differentiated adenocarcinoma

### Differential gene expression in GBC compared with the control samples

Analysis of 13 GBC and 6 control samples yielded an average of 40.64±4.3 and 39.41±4.73 million reads, respectively, with 70% mapping rate to the human reference genome. The basic read quality and alignment statistics have been summarized in supplementary figure S5. Principal component analysis (PCA) showed PC1 accounting for 28% and PC2 for 22% of expression variance among samples (figure 1a). Differential expression analysis with 16,231 transcripts qualifying for analysis (≥10 read count in at least one sample) identified 1,319 genes to be differentially expressed-528 (∼40%) upregulated and 791 (∼60%) downregulated. Volcano plot analysis revealed HOXB7 and COL11A1 as highly expressed in GBC, while FGF19 and CHST4 were the most downregulated genes(figure 1b). Top differentially expressed genes (DEGs) have been listed in supplementary table S2.

**Figure 1:**
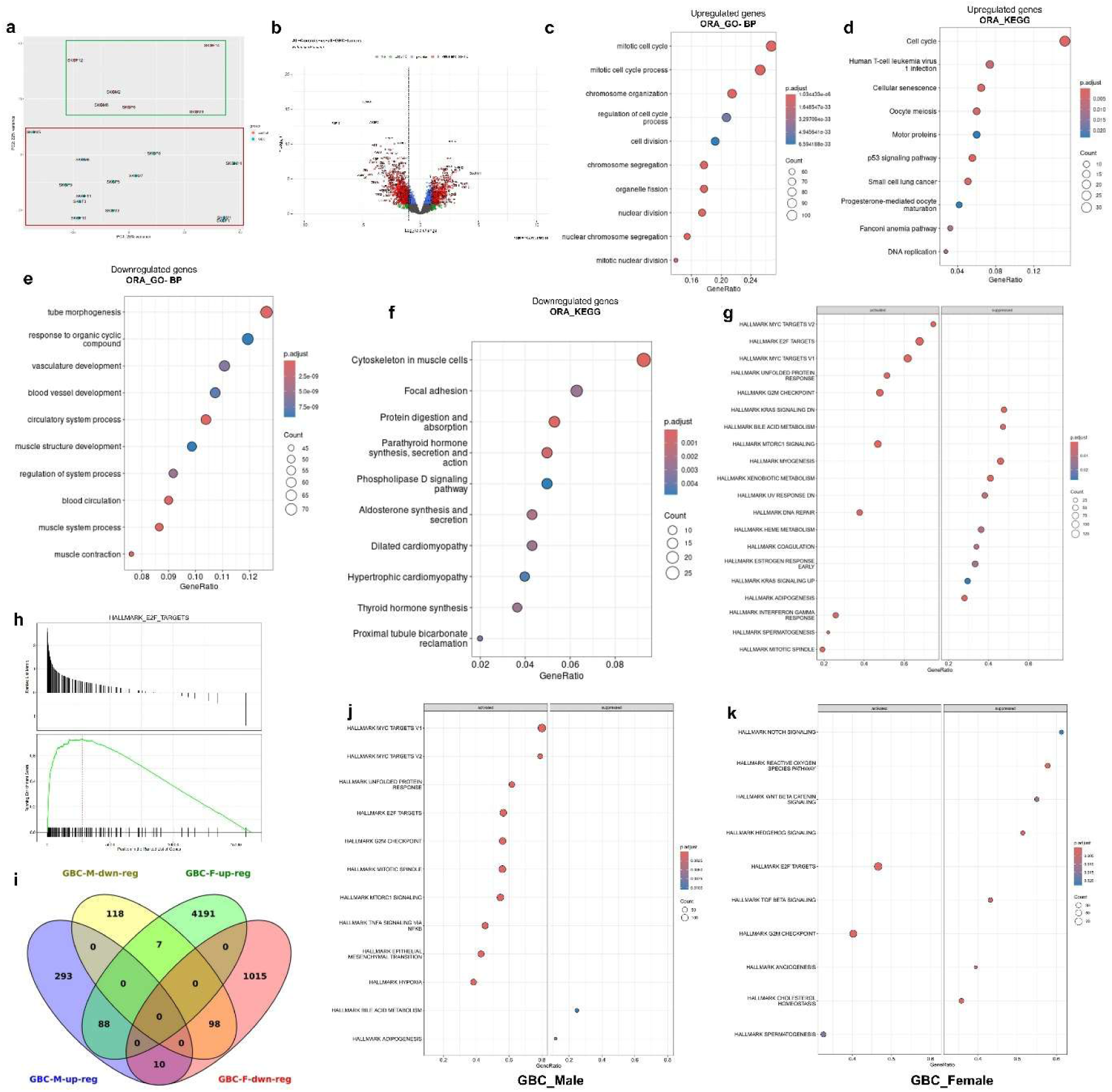
Analysis of RNA-seq data. a PCA plot showing clustering between GBC case and control samples. b.Volcano plot showing expression of different genes. X axis denotes the log2(fold change) of genes while the Y axis denotes the statistical significance. The differentially expressed genes are marked as red dots. c,d. Over representation analysis using upregulated genes with Gene Ontology-Biological Process database and KEGG database. e,f Over representation analysis using downregulated genes with Gene Ontology-Biological Process database and KEGG database. g GSEA bubble plot showing different pathways either activated or suppressed in gallbladder cancer (MSigDB). x-axis represents the gene ratio (proportion of differentially expressed genes contributing to each pathway), bubble size indicates the number of genes contributing to the pathway and the color gradient indicates the statistical significance. h GSEA of one of the top activated pathway Hallmark_E2F_Targets. i Venn diagram showing common and exclusive upregulated or downregulated genes in Male and Female GBC. j,k GSEA bubble plot showing different pathways either activated or suppressed in gallbladder cancer (MSigDB) of male (j) or female (k).

Gene Ontology (GO) analysis of the 528 upregulated genes with showed significant enrichment in 338 biological processes (GO-BP) terms (figure 1c) (e.g., mitotic cell cycle, chromosome organization), 23 molecular functions (GO-MF) terms (e.g., microtubule binding, kinase activity), and 79 cellular components (GO-CC) terms (e.g., centromeric regions, mitotic spindle) (figure S6a, S6b). KEGG pathway analysis with the upregulated genes highlighted the cell cycle and p53 signaling pathways as key in GBC (figure 1d). GO analysis of 791 downregulated genes revealed enrichment in 490 biological processes terms (e.g., circulatory system, muscle function) (figure 1e), 37 molecular functions terms (e.g., extracellular matrix binding), and 87 cellular components terms (e.g., extracellular matrix, cell surface) (figure S6c, S6d). KEGG pathway analysis with the downregulated genes included pathways related to cytoskeleton, cell adhesion molecules etc (figure 1f).

GSEA identified activated pathways such as E2F targets, G2/M checkpoint, MYC targets, and DNA repair, indicating increased proliferation and genomic instability. Suppressed pathways included bile acid metabolism, KRAS signaling, and xenobiotic metabolism, suggesting disrupted metabolic regulation (figure 1g and S7a,b). E2F targets were of highest enrichment score (NES = 0.73) (figure 1h), highlighting their key role in GBC progression and poor prognosis. The intricate interactions among E2F factors, cyclins, CDKs, and the Rb protein form a complex regulatory network that guarantees accurate control of the cell cycle (8). Its association with tumorigenesis and poor prognosis has been well documented in the literature (9,10). GSEA using GO-BP and KEGG database highlighted pathways related to DNA replication and cell division (figure S8a,b).

To validate the RNA-seq findings, we performed qPCR for six differentially expressed genes. Downregulation of FGF19, CHST4, and AMBP and upregulation of COL11A1, HOXB7 and KRT17 in GBC were confirmed by qPCR (figure S9).

### Differential gene expression between male and female cases of GBC

The comparative analysis of DEGs in male and female GBC cases was performed to identify sex specific, distinctive patterns in the gene expression. We identified a total of 88 upregulated and 98 downregulated genes common in both the sexes. Further, 293 genes were uniquely upregulated in male GBC, while 4191 genes were unique to female GBC. Among the downregulated genes, 118 were unique to male GBC, and 1015 were unique to female GBC (figure 1i). Interestingly, we also observed 17 genes that showed opposite trends in female and male cases (Table 2).

**Table 2:** Differentially Expressed Genes with opposite trends in male and female GBC.

| Upregulated in male GBC downregulated in female |  | Upregulated in female GBC downregulated in male |  |
| --- | --- | --- | --- |
| GATA5 | PGGHG | MALRD1 | HOXA5 |
| REG1A | TYPR1 | LEP | LOC101927057 |
| LCN2 | UNC5B | ABCB5 |  |
| ST6GALNAC1 | STC2 | KCNIP2 |  |
| LTBP1 | LOC105369230 | ACVR1C |  |

Gene set enrichment analysis revealed pathways with similar dysregulation in males and females as well as few unique one (figure 1j & 1k). K-ras signaling, a pathway crucial for cell growth and survival, was activated as evident by the downregulation of set of genes which are downregulated by k-ras. Bile acid metabolism, which is a characteristic feature of biliary epithelium, was downregulated in GBC irrespective of the sex. This indicates the dedifferentiation of epithelial cells, a hallmark of cancer. A few other pathways, like E2F targets, MYC targets, and G2M checkpoints, were upregulated both in male and female GBC samples. Interestingly, multiple pathways showed significant but different trends in male and female cases of GBC.

### Differential Gene Expression in Male GBC Compared to Male Control GB

When male GBC samples (n=6) were compared with unmatched control GB tissues (n=3), a number of DEGs were identified. The top gene list is described in table 3. A more elaborated list of DEGs may be found in supplementary table S3.

**Table 3:** Top differentially expressed genes in Male GBC.

| Downregulated |  | Upregulated |  |
| --- | --- | --- | --- |
| PLIN1 | FGF19 | CSCL3 | ZNF865 |
| UGT2B28 | UGT2B15 | IGFBP3 | CEMIP |
| FABP4 | ASGR2 | ANXA8L1 | KRT16 |
| PPP1 | PLIN4 | ESM1 | CILP2 |
| R1A | CD36 | CCL20 | COL11A1 |
| CES1 | LPL | INHBA | MUC2 |
| ASGR2 | XIST | BPIFB1 | HSPA5 |

The analysis showed that several genes involved in lipid metabolism, such as PLIN1, FABP4, and UGT2B28 were significantly downregulated in male GBC. Fatty Acid Binding Protein 4 (FABP4), also known as adipocyte FABP (A-FABP) facilitates fatty acid transport and can alter lipid metabolism in cancer cells (11). GBC often exhibits dysregulated lipid metabolism that can contribute to proliferation and survival, possibly by FABP4 downregulation (12). Additionally, FGF19, which regulates bile acid production and metabolism, was also downregulated. In contrast, genes associated with extracellular matrix remodeling, including CEMIP, COL11A1, CCL20, INHBA, and ESM1, were prominently upregulated. These proteins can contribute to processes like matrix degradation and cell migration, which are critical for tumor progression.

### Differential Gene Expression in Female GBC Compared to Female Control GB

When female GBC tissues (n=7) were compared with unmatched control GB (n=3), patterns of differential gene expression were obtained. The top gene list is described in table 4. A more elaborated list of DEGs may be found in supplementary table S4.

**Table 4:** Top differentially expressed genes in Female GBC.

| Downregulated |  | Upregulated |  |
| --- | --- | --- | --- |
| ACTG2 | VTN | GRIN2B | HOXB7 |
| SYNM | LGALS2 | CENPF | CENPE |
| AGT | DES | SOX11 | SLC6A14 |
| FAM107A | ASGR2 | WDR87 | MUC4 |
| NR2F1 | CNN1 | FAM133A | ASPM |
| PI16 | SLC9A3 | SERPINB5 |  |
| AMBP | GC | IGF2BP3 |  |
| UGT2B15 |  |  |  |

Genes associated with cytoskeletal integrity, such as ACTG2, SYNM, and CNN1 were significantly downregulated in females, reflecting cytoskeletal disorganization and lack of structural integrity. In addition, NR2F1, a transcription factor known to regulate differentiation (13), was also downregulated in female GBC samples. Contrary to this, VTN (vitronectin) and MUC4 (mucin 4) were upregulated. Overexpression of these proteins is consistent with their roles in extracellular matrix remodelling and tumor progression (14,15). HOXB7, a transcription factor that is known to promote angiogenesis, is also upregulated in cancer tissues (16).

### Exclusive Hallmark Pathways in male GBC

Multiple Hallmark immune pathways were significantly upregulated in male GBC cases. These included Interferon Alpha and Gamma Response, Hallmark TNF Alpha Signaling, Hallmark IL2 STAT5 Signaling (supplementary figure S10a), Hallmark IL6 JAK STAT3 Signaling (supplementary figure S10b) and Hallmark Interferon Gamma response (supplementary figure S10c). This highlights the involvement of immune and inflammatory responses specifically in male GBC tumor progression.

Additionally, both androgen and estrogen response pathways were found to be upregulated in male GBC (supplementary figure S10d), suggesting a role of sex hormones in the pathogenesis of GBC. Surprisingly, no similar pathway was significantly dysregulated in female GBC.

### Exclusive Hallmark Pathways in female GBC

In females, Hedgehog signaling was observed to be downregulated significantly (supplementary figure S11a). Hedgehog signaling is known to play an essential role in tissue homeostasis, and aberrant Hedgehog signaling has been associated with various cancers (17). Interestingly, studies are showing that estrogen receptor plays a role in Hedgehog signaling (18), suggesting an interplay between these two pathways and downregulation of hedgehog signaling, specifically in female GBC.

In addition, metabolic pathways like cholesterol homeostasis have been downregulated in female GBC cases. The downregulation of cholesterol homeostasis suggests there may be an altered lipid metabolism, which has been recognized as a hallmark of cancer progression. Bile salts and bile acids have been implicated in gallbladder cancer (GBC) development, which could occur from dysregulated cholesterol synthesis (19) as evident by deranged bile acid metabolism (supplementary figure S11b).

### Construction of co-expression network and identification of the modules

A total of 16,231 annotated transcripts were selected for differential gene expression analysis, and all DEGs were used to construct the co-expression network. The optimum soft-thresholding power β was chosen based on the correlation coefficient between log(ϰ) and log[p(ϰ)] under different values of the soft-thresholding power, β. The β was selected as 8 to ensure the scale-free topology (figure 2a,b). When the β was 8, the scale-free topology (R^2^) was 0.85 (figure 2c,d). After selecting the soft-thresholding power β, the gene expression profile of 16,231 genes was transformed into an adjacency matrix, and TOM and disTOM were subsequently constructed. The co-expression network was then built. The identification of modules was achieved using a dynamic tree cut method with a minimum size of 30 (figure 2e,f). All 16,231 genes were segregated into different clusters, while no gene was assigned to the grey module. A total of 115 modules were initially identified, and after merging similar modules using a merging threshold function of 0.25, we obtained 27 modules with co-expressed genes. The mean expression values (MEs) of the merged modules were then calculated.

**Figure 2:**
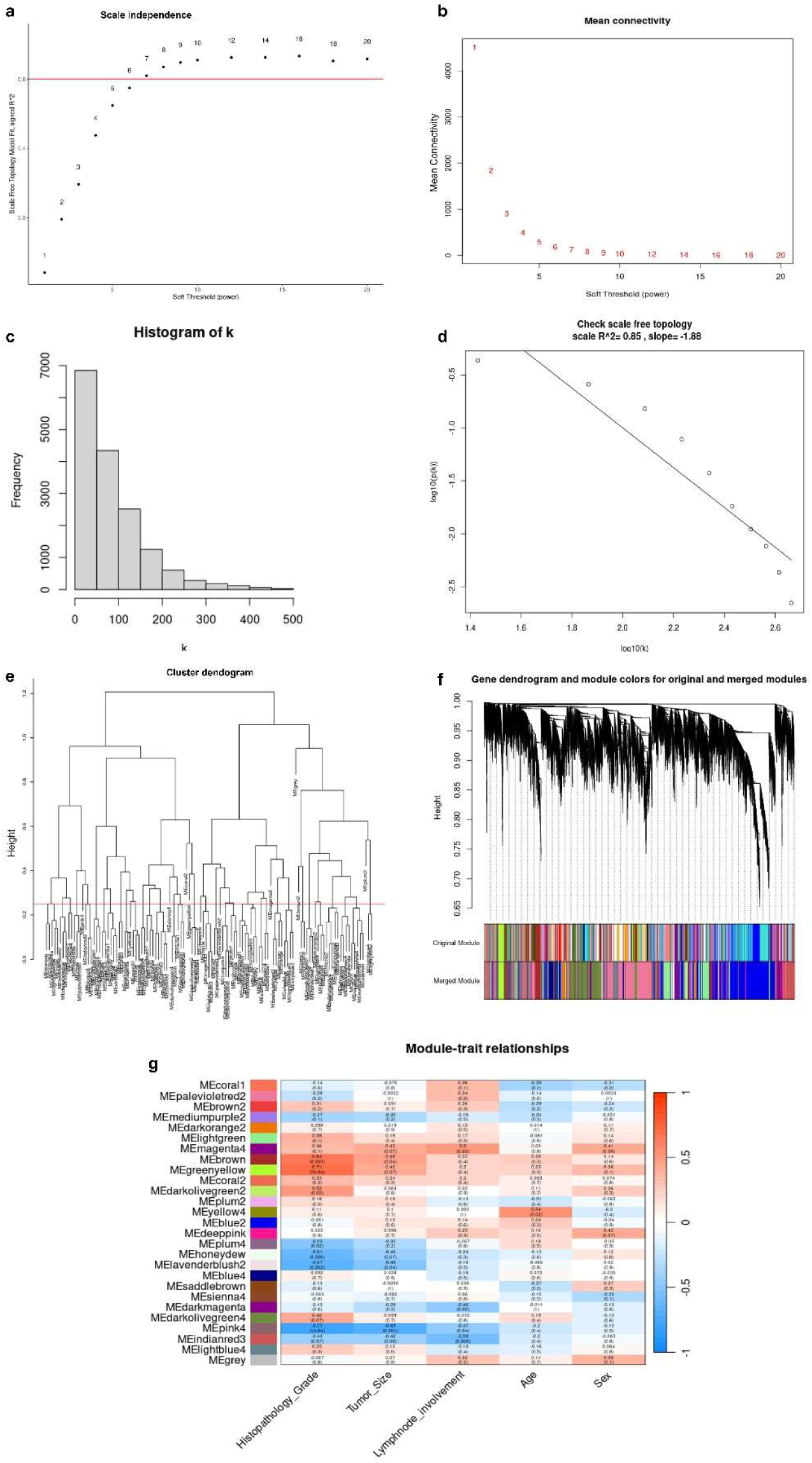
Construction of WGCNA. a,b Analyses of network topology of various soft-thresholding powers (weighting coefficient β). The x-axis represents different soft-thresholding powers. The y-axis represents the correlation-coefficient between log(k) and log[p(k)], was set at 0.8. c,d Histogram of connectivity distribution and scale free topology (R2=0.85) when β=8. e,f Clustering dendogram of the genes involved. g Heatmap of the correlations between MEs and the clinical traits. Correlation coefficients and corresponding p values are shown in the rectangles and the brackets respectively.

### Gene co-expression modules associated with clinical traits

The Pearson correlation coefficients were determined for the MEs of all 27 merged modules with the clinical trait information for all 19 samples to investigate the module-trait relationship (Figure 2g). According to the module-trait relationships, the green-yellow (*r* = 0.71, *p* = 7e-04) and brown modules (*r* = 0.64, *p* = 0.003) have shown the strongest association with the clinical trait - histopathology grade. Further, to check how the clinical trait histopathology grade fits with the green-yellow and brown modules, a hierarchical clustering dendrogram and a heat map were constructed, revealing that the green-yellow and brown modules were tightly correlated with the histopathology grade (figure 3a). Further, the GS for histopathology grade was calculated for each module to ensure the reliability of the selection of significant modules for the clinical trait of concern. The mean absolute values of GS in each module were calculated and visualized, suggesting that the brown module was the highest clinically significant module with the highest GS for histopathology grade (figure 3b).

Furthermore, the MM values were obtained, and the correlation analysis was performed between the GS of histopathology grade and the MM for genes in the green-yellow and the brown module. Based on the expression and correlation analysis of the two modules - green-yellow (*r* = 0.5, *p* = 2e-18) (figure 3c,e) and brown (*r* = 0.54, *p* = 2.8e-42) (figure 3d,f), the brown module was selected as the clinically significant module for GBC in terms of histopathology grade. Thus, we focused on the brown module for further investigation of biological functions, pathways analysis and identification of hub genes (supplementary figure S12). The complete gene list of Brown module is mentioned in supplementary table S5.

**Figure 3:**
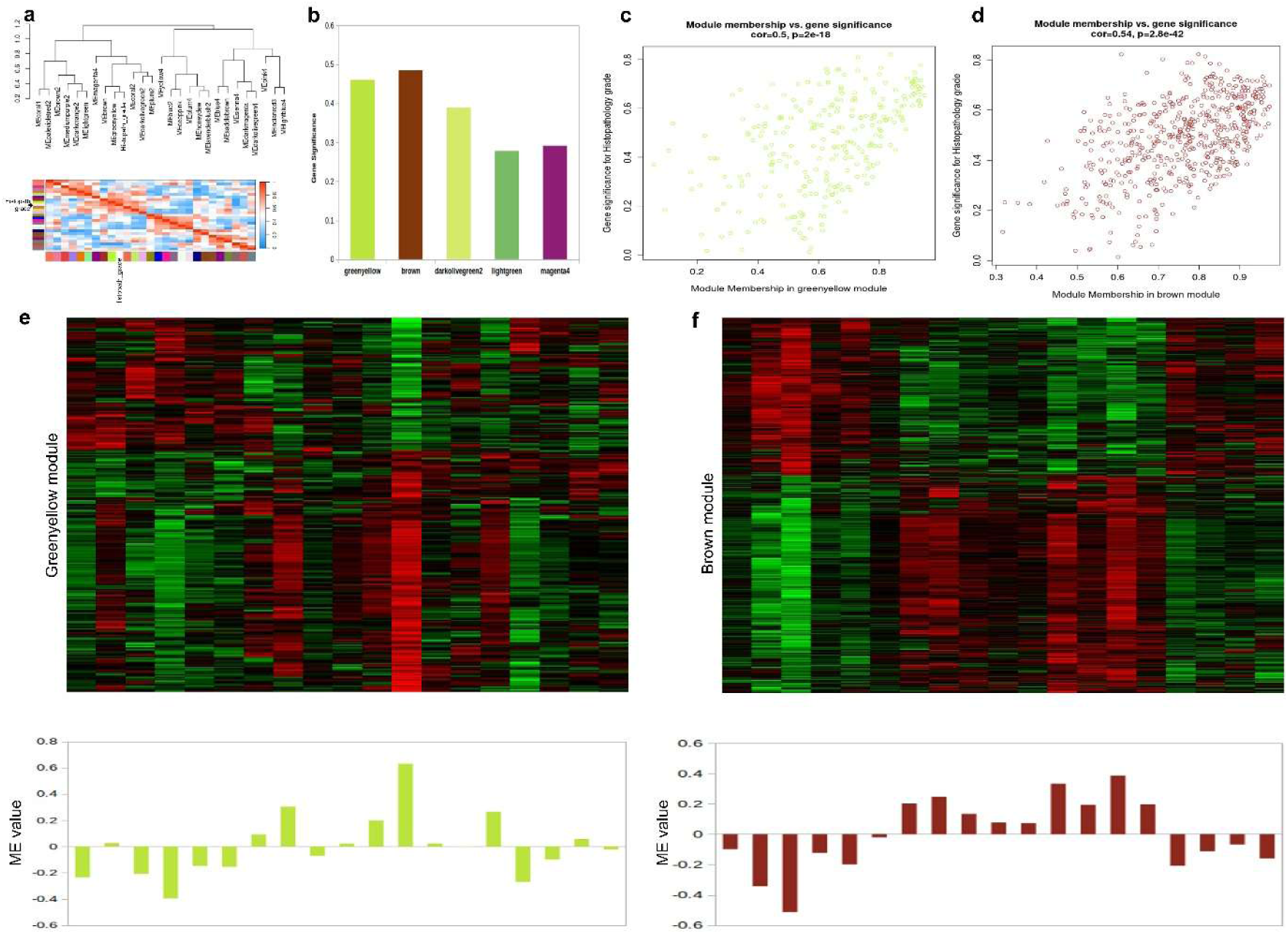
Identification of clinically significant modules. a Hierarchical clustering dendogram of MEs and Histopathology grade. Heatmap plot shows the adjacencies among them. Red color indicates positive correlation, while blue color indicates negative correlation. b Bar plot of mean absolute values of GS across modules. Higher mean GS suggests more significant associations between module and the histopathology grade. c,d Scatterplot of GS for histopathology grade versus MM in the greenyellow and the brown module. e,f The upper heat maps show the expression levels of all genes in the greenyellow and the brown module (y-axis) among all samples (x-axis). The lower bar diagrams show the corresponding MEs(y-axis) versus the samples (x-axis).

### Functional enrichment and pathway analysis and hub genes identification of the brown module

Based on the filtering criteria of MM > 0.6 and q.weighted < 0.001 that were applied on the 541 genes from the brown module, we finally reached the 110 filtered genes (Additional file 2, Table S6), where the gene ontology (GO), functional enrichment and KEGG pathway analysis was performed. The top 10 significantly enriched terms of biological process (BP) were identified, suggesting that the brown module genes are mainly associated with mitotic cell cycle processes (figure 4a). KEGG pathway analysis of 110 genes indicates that the genes are critically involved in the cell cycle and p53 signaling pathways (figure 4b). To identify the hub genes in the brown module, the top 20 genes with the highest IC were extracted to construct the gene-gene interaction key network, considered as central genes. The connections among central genes with each other were visualized (figure 4c).

**Figure 4:**
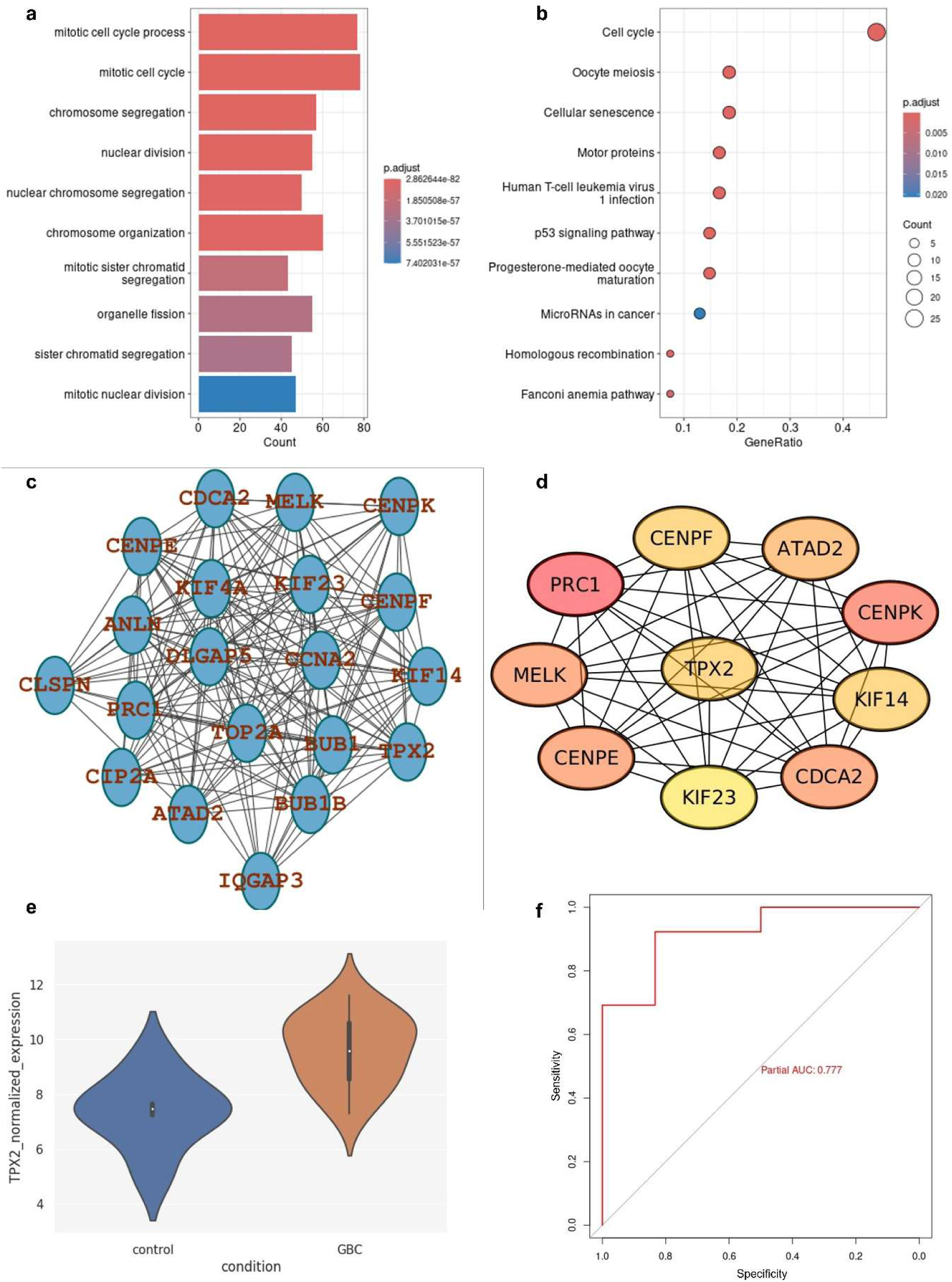
Functional enrichment analysis and hub genes prediction of the brown module. a Bar chart of the top 10 significant terms of BP in GO enrichment analysis of the brown module. b Dot plot of the top 10 significantly enriched KEGG pathways of the brown module. c Interaction network in the brown module. The top 20 genes with the highest IC were associated. d Top 10 hub gene prediction using DMNC algoritm from Cytohubba module. e Contour plots of TPX2 expression levels between control and GBC group. f ROC analysis to evaluate expression level of TPX2 in predicting GBC.

Finally, TPX2 was selected as the central hub gene of the clinically significant module as it had the highest degree of connections among 10 genes (figure 4d). Further, we focused on the key hub gene TPX2 for additional analysis. The expression of TPX2 was significantly upregulated in GBC compared to the control samples (figure 4e). In addition, ROC analysis revealed that TPX2 could positively predict the diagnosis of GBC (AUC = 0.69) (figure 4f).

### Survival analysis

We further validated the top 12 overexpressed genes identified by WGCNA based on their survival information and expression pattern using the cholangiocarcinoma (CHOL) cancer type in TCGA database. All the hub genes were found to be either significantly overexpressed or showing a prominent elevated pattern in the database (figure 5a-d, supplementary figure S13). Kaplan-Meier curve analysis was conducted in patients with cholangiocarcinoma (CHOL in TCGA) to investigate overall survival in association with the top hub genes. One of the central hub genes KIF23, was found to be associated with poor survival with a Hazard ratio of 2.2 and p p-value 0.12. Most of the other hub genes, like BUB1B, TOP2A, CLSPN, etc showed a similar trend (although a non-significant p value that might be attributed to the smaller number of cases for CHOL in the TCGA database). Nonetheless, we found strong data coherence with the publicly available database.

### Validation of selected hub genes

Hub gene expression strongly correlated with GBC histopathological grade, with qPCR validation confirming significantly higher expression in GBC versus controls (figure 5e-j). All hub genes exhibited increasing trends with higher tumor grades, although statistical significance was limited by the small sample sizes in some groups for the groups (WDADC and MDADC).

Targeting Protein for Xenopus kinesin-like protein 2 (TPX2) emerged as a key hub gene with significant upregulation in GBC tissues in graded manner. it is a microtubule-associated protein essential for mitotic spindle assembly and chromosomal stability. TPX2 activates and stabilizes Aurora kinase A (AURKA) which is known to promote cell cycle, thus forming a potent oncogenic axis (20). This interaction promotes unchecked cell cycle progression, centrosome amplification, and genomic instability, driving tumor initiation and progression. Expression of TPX2 was verified to be increased in GBC compared to control GB tissue by immunohistochemistry (figure 5k).

**Figure 5:**
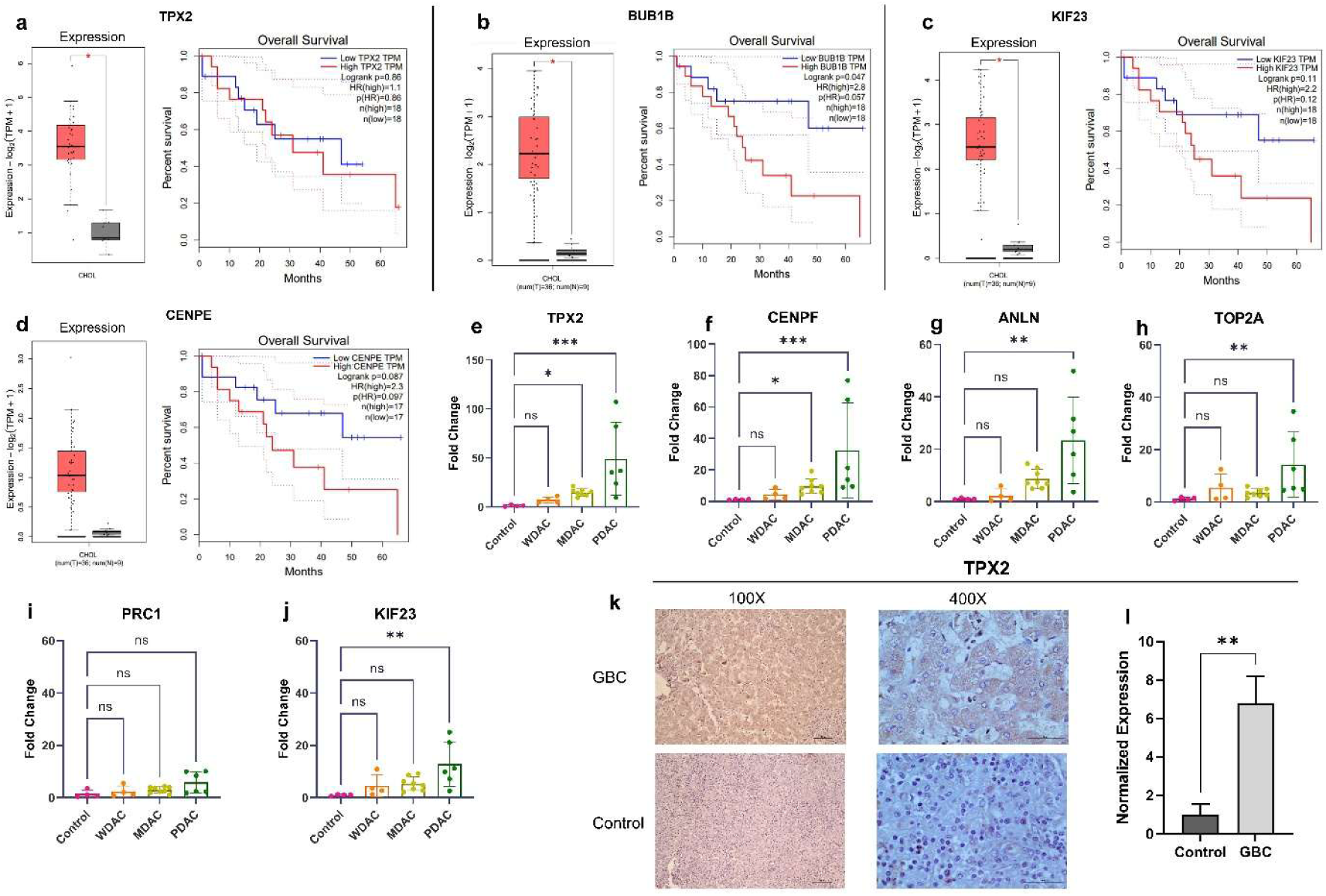
Verification of hub genes. a-d Kaplan Meier survival analysis based on expression of different hub genes in cholangiocarcinoma (CHOL) type cancer (TCGA database). Red boxes represent cholangiocarcinoma while grey boxes indicate control. e-j Expression of selected hub genes across different grades of GBC tissues. Fold change of the target gene is with respect to the expression in control normalized with housekeeping gene GAPDH. k,l Expression of TPX2 in GBC and control tissues. The bar diagram on the right shows the normalized expression of the target. *p value <0.05, **p value <0.01, ***p value <0.001.

### Aurora Kinase inhibitors result in attenuation of cancer phenotype in vitro

Overexpression of AURKA in GBC tissue was verified by immunohistochemistry (figure 6a). We then treated invasive GBC cell line G415 with TC-S7010 (inhibitor of Aurora kinase A) and hesperadin hydrochloride (inhibitor of Aurora kinase B) and checked for different components of cancer phenotype e.g. proliferation, cell migration (by wound healing assay) as well as molecular markers for EMT and invasion (MMPs and TIMPs). Our results showed significant downregulation of EMT supported by the downregulation of mesenchymal markers like N-cadherin and vimentin along with upregulation of epithelial marker E-cadherin in the presence of Aurora kinase inhibitors (figure 6c-e). We also observed downregulation in the MMPs (MMP2 and MMP9), along with variable modulation of TIMPs (TIMP1 and TIMP2) in the presence of these inhibitors (figure 6f-i). Furthermore, there was reduction of cell migration as well evident by delayed wound healing following treatment with (figure 6j,k). Additionally, treatment with the inhibitors resulted in significant increase in G0/G1 phase (nuclear content 2n) of cell cycle as well as significant reduction in S phase and G2/M phase (nuclear content 4n) (Additional file 1, figure S14). All these findings imply that Aurora kinase inhibitors attenuate cellular proliferation and cancer phenotype.

**Figure 6:**
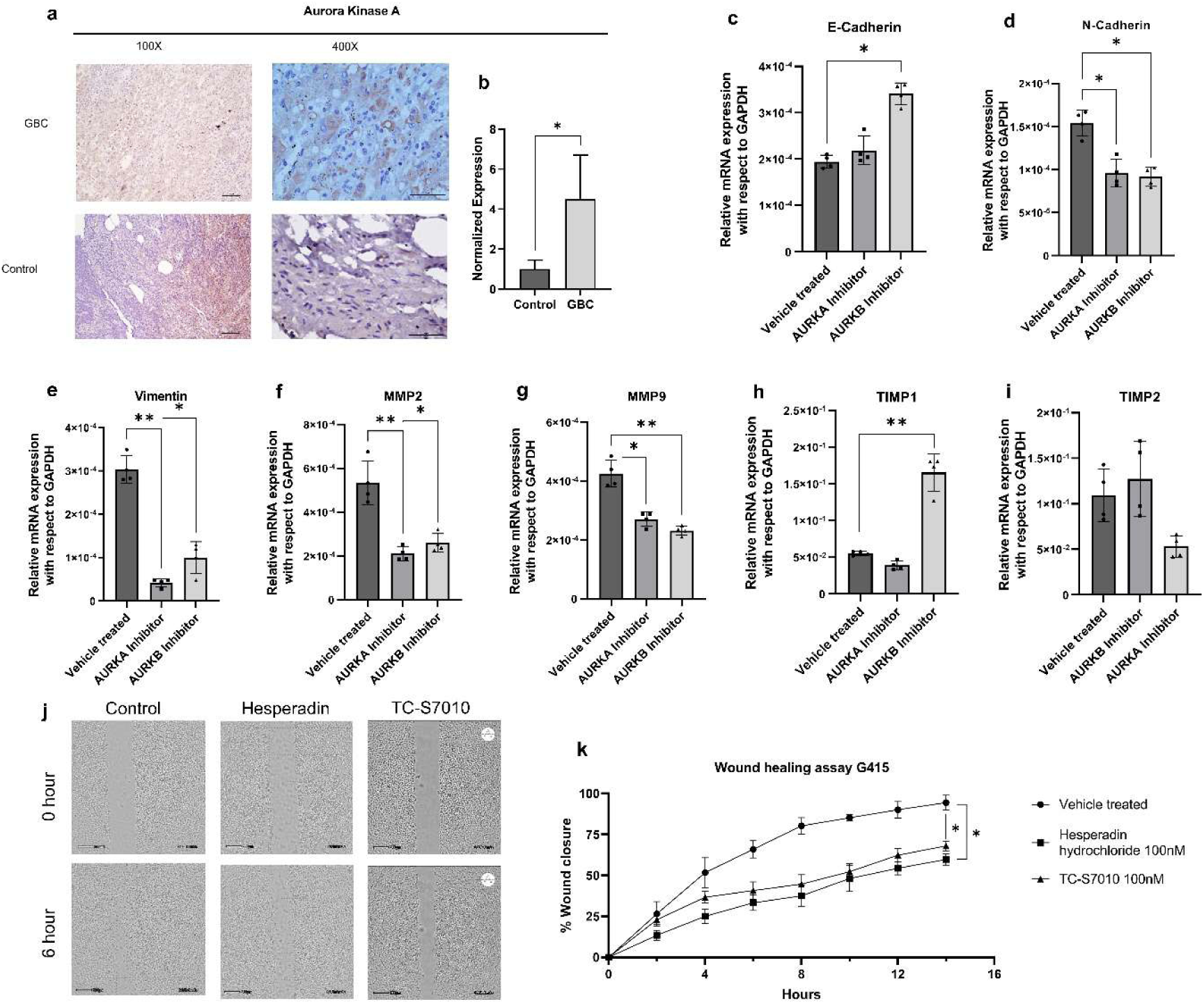
Effect of Aurora Kinase inhibitors on GBC cell line G415. a,b Expression of AURKA in GBC and control tissues. Bar diagram on the right shows the normalized expression of the target c-e Expression of E-cadherin was upregulated along with downregulation of N-cadherin and vimentin. f-i. Downregulation of both MMP2 and MMP9 was observed. Effect on expression of TIMP1 and TIMP2 was variable. All the expression values are relative to GAPDH. j,k Treatment with hesperadin hydrochloride an inhibitor of AURKB and with TC-S7010 showed significant reduction in cellular migration and delayed wound healing in gallbladder cancer cell line G415. *p value <0.05, **p value <0.01.

## Discussion

Our study provides a comprehensive transcriptomic analysis of GBC using RNA sequencing, revealing differentially expressed key genes (DEGs) and deranged biological pathways. By means of WGCNA, we also identified highly interconnected hub genes closely associated with the histopathological grades of GBC. The demographic distribution aligns with previous epidemiological studies (21), which have shown a higher incidence of GBC in elderly populations. Moderately differentiated adenocarcinoma (MDADC) is predominant, with most cases being at advanced stages (Stage III and IV) at the time of diagnosis, emphasizing the aggressive nature of this tumor and the need for strategies for early diagnosis. We further demonstrated that there is significant heterogeneity within the tumor group, indicating a complex molecular landscape of GBC.

Dysregulation of bile flow may cause chronic inflammation leading to neoplastic changes of the biliary tract. Fibroblast growth factor-19 (FGF19), highly expressed in gallbladder epithelial cells, maintains bile acid homeostasis by inhibiting hepatic synthesis and secretion through a negative feedback loop (22). Downregulation of FGF19 signifies dedifferentiation of GB epithelium as well as higher bile acid output from the liver. These may lead to chronic inflammation and altered cell growth (23).

CHST4 (Carbohydrate Sulfotransferase 4), another protein found to be downregulated in gallbladder cancer, encodes an enzyme that causes the sulfation of carbohydrates, specifically mucin-associated glycans. These glycans play a crucial role in lymphocyte homing ─ a process that directs immune cells to specific tissues and locations, such as inflammation sites (24). Recent studies have shown that CHST4 is abnormally expressed in various types of cancers and can act as a potent biomarker. A study by Zhang et al. showed that CHST4 is downregulated in hepatitis B virus-associated hepatocellular carcinoma tissues compared to normal liver tissues and this downregulation correlates with increased viral replication as well as tumor progression (25). Conversely, higher expression of CHST4 was associated with tumor suppression and the creation of an effective immune response. Gallbladder cancers often create an inflammatory microenvironment where CHST4 might play a role in maintaining immune homeostasis. This signifies the potential role of CHST4 as a tumor suppressor and a therapeutic agent (26).

Our study revealed significant upregulation of COL11A1, HOXB7, and KRT17 in gallbladder cancer tissues. COL11A1, a type XI collagen component crucial for extracellular matrix remodeling, is upregulated across multiple cancers including breast (27), ovarian (28), pancreatic (29), colorectal (30), and lung cancer (31). It activates cancer-associated fibroblasts, enhancing cancer cell migration through interactions with integrin α1β1 and DDR2 receptors, triggering TGF-β1 and NF-YA signaling cascades that promote invasion (32). COL11A1 drives cancer progression by creating dense fibrotic stroma that facilitates invasion, impedes drug delivery, and confers chemoresistance. High expression correlates with poor prognosis and serves as a potential biomarker for differentiating pancreatic adenocarcinoma from other conditions (33).

We also found an upregulation of HOXB7 in GBC tissues, consistent with previous upregulation reported in hepatocellular carcinoma, where overexpression in HOXB7 has been shown to increase the expression of MMP2 and MMP9, enzymes that break down extracellular matrix components and have been known to promote tumor cell invasion (34). HOXB7 has also correlated with poor prognosis and disease aggression in cholangiocarcinoma supporting its potential as a biomarker (35). In another study, HOXB7 has been found to promote angiogenesis through increased expression of angiogenic factors VEGF and IL-8 (36). Other studies have demonstrated that HOXB7 increases metastasis in intrahepatic cholangiocarcinoma (ICC) and liver cancer (37). Our results support these findings, suggesting that HOXB7 may be contributing to the progression and metastatic potential of GBC, possibly through similar mechanisms.

KRT17, a cytokeratin intermediate filament protein and epithelial cell marker, showed significant upregulation in GBC versus controls. This aligns with KRT17’s established role in gastric, thyroid, and esophageal cancers. Previous studies demonstrated KRT17 overexpression in lymph node-positive GBC and its potential as a marker of micro-metastatic (38). KRT17 knockdown inhibits lung cancer cell invasion and proliferation, while overexpression activates Wnt signaling and downstream targets (cyclin D1, c-Myc, MMP7), confirming the association between cytokeratins and metastasis (39).

Our data shows the activation of mTORC1 signaling pathway in GBC. The mammalian target of rapamycin (mTOR) pathway regulates cell proliferation, growth, and survival. Dysregulation of mTOR has been documented in a variety of cancers (40). Interestingly, KRT17 induces activation of the mTOR pathway, while its knockdown reduces the phosphorylation of mTOR and S6K1, its downstream effector (41). Overall, KRT17, along with HOXB7, may drive proliferation and invasion through the mTOR/S6K1 axis in GBC.

Gene Set Enrichment Analysis (GSEA) revealed multiple dysregulated pathways in GBC. Key upregulated pathways included E2F targets, which regulate cell cycle progression, DNA replication, and proliferation, frequently contributing to tumorigenesis (42). The G2M checkpoint pathway, controlling cell cycle progression from G2 to mitotic phase, showed upregulation indicating dysregulated cell cycle control enabling uncontrolled cancer cell proliferation (43). MYC targets, under control of the proto-oncogene MYC that regulates cell growth and differentiation, were also upregulated, signifying enhanced cell proliferation and survival characteristic of cancer (44). Unfolded protein response (UPR)stimulates the expression of pro-angiogenic factors in response to hypoxia, thereby promoting angiogenesis. UPR-mediated rewiring of metabolic pathways also allows the tumour to cope with nutrient deprivation (45). One mechanism by which cells can maintain UPR activation while avoiding initiation of cell death is actively suppressing components of the cell death pathway or stimulating cytoprotective mechanisms such as autophagy (46).

Each pathway plays a crucial role in regulating various aspects of the cell cycle and cellular functions associated with cancer development. The significant upregulation of these pathways in the tumor samples underscores their importance in driving GBC.

Sex specific analysis identified exclusive dysregulated genes and pathways in male and female cases of GBC. Both perilipin 1 and 4, essential lipid droplet surface proteins regulating energy balance and lipid metabolism, were downregulated in male GBC. PLIN1 downregulation has been reported in hepatocellular carcinoma, where overexpression reduced cell proliferation, migration, and invasion (47). PLIN4, primarily found in white adipose tissue and skeletal muscle, shows dysregulation in gastric, lung, and basal-like breast cancers with prognostic implications (48–50).

SOX11, one of the top upregulated genes in female GBC, has been implicated across multiple cancer types in pan-cancer analyses, correlating with survival outcomes, stage, and recurrence. SOX11 expression negatively correlates with CD8+ T cell infiltration, particularly in colon, pancreatic, kidney, lung, and thyroid cancers (51), suggesting immune-related mechanisms in female GBC progression.

WGCNA analysis identified a network of highly connected genes associated with GBC pathological grade, including hub genes TPX2 and KIF23. KIF23, a kinesin-like motor protein involved in microtubule-dependent organelle transport and chromosome movement during cell division, has been established as an independent prognostic factor in lung cancer and implicated in hepatocellular (52), endometrial, esophageal (53), and triple-negative breast cancers (54).

TPX2 (Targeting Protein for Xklp2) is amplified in genomically unstable cancers, including gastric, colon, oral squamous cell, and ovarian cancers. Initially identified as a microtubule-associated protein directing Xklp2 localization, TPX2 also binds, activates, and localizes Aurora A kinase (AURKA) to the spindle during mitosis (55). AURKA plays crucial roles in centrosome maturation and spindle assembly, with Aurora kinases classified as a family of oncogenic proteins (56). Its functional interaction with Aurora kinases forms a potent oncogenic axis that promotes cell cycle progression, centrosome amplification, and genomic instability (57).Independent validation across multiple patient cohorts confirmed Aurora kinase overexpression in advanced-stage tumors, correlating with reduced overall survival (58,59). Co-expression and co-amplification patterns of TPX2 and Aurora kinases underscore their collaborative role in GBC pathogenesis through aberrant mitosis and chromosomal missegregation.

In a separate approach we performed screening of 210 kinase inhibitor compounds in G415 gallbladder adenocarcinoma cells (data not shown) and identified TC-S-7010 (an Aurora A inhibitor) and Hesperadin hydrochloride (an Aurora B inhibitor) as having the most potent anti-proliferative effects. Immunohistochemistry confirmed enhanced Aurora kinase expression in high-grade GBC patient tissues, supporting its therapeutic potential.

Aurora kinase inhibition profoundly impacts epithelial-mesenchymal transition (EMT), reducing mesenchymal markers (N-cadherin, vimentin) while upregulating epithelial marker E-cadherin. Decreased MMP2/MMP9 and modulated TIMP1/TIMP2 expression, along with delayed wound closure following Hesperadin hydrochloride treatment, confirmed suppression of invasive and metastatic potential.

These findings establish a mechanistic link between the TPX2-Aurora kinase axis and EMT regulation in GBC, positioning Aurora kinase inhibitors as promising therapeutic strategies to halt disease progression and metastasis in this aggressive malignancy with limited treatment options.

## Conclusions

TPX2 and Aurora kinases, key cell division regulators frequently dysregulated in cancer, govern mitotic spindle assembly and chromosome segregation. WGCNA analysis and experimental validation identified TPX2 as a central hub gene associated with GBC tumor histopathological grade, likely through TPX2-mediated AURKA activation, promoting increased proliferation and higher tumor grades. These findings suggest that Aurora kinase inhibitors may be promising therapeutic strategies for GBC, although validation in larger cohorts is needed.

Our data reveal a complex, heterogeneous molecular landscape in GBC, with cell cycle regulation, DNA replication, and Aurora kinase signaling as major tumorigenic drivers. Gender-specific differences emerged: male GBC samples showed altered E2F pathway activity, while female samples demonstrated cytoskeletal dynamic changes, suggesting sex-specific disease mechanisms that require further mechanistic studies for early diagnosis and targeted therapy implications.

While offering valuable insights into GBC molecular underpinnings, our study has the limitation of a relatively small sample size, which limits statistical robustness. Future studies with larger, diverse cohorts will be critical to validate and extend these observations.

## Acknowledgements

S.K. and R.D. want to express their sincere gratitude to the Department of Biochemistry, AIIMS, New Delhi for providing infrastructure and space to carry out this research.

## Authors’ contributions

S.K.S. performed the experiments with assistance from R.M., G.S., N.B., D.K., under the supervision of R.D. and S.K.. A.N. helped with bioinformatics analysis. P.D. helped with the histopathological assessment of tissue samples. G.S. and D.K. performed validation experiments. N.D. and K.K. performed the surgical procedure. R.D. assisted in IHC assays and quantitation. S.K. and R.D. oversaw the entire work.

## Ethics approval and consent to participate

The study was approved by Institute Ethics Committee of AIIMS, New Delhi. Informed consents were obtained from all the participants. The study was conducted in accordance with the Declarations of Helsinki.

## Consent for publication

All the authors consented for publication.

## Data Availability

RNA sequencing data were deposited into the Bio Project (NCBI) with accession link https://www.ncbi.nlm.nih.gov/bioproject/PRJNA1242580. The original contributions data presented in the current study are included in the manuscript/ supplementary material; further inquiries are available from the corresponding author upon reasonable request.

## Competing interests

The authors declare no competing interests.

## Funding information

This work was supported by the Department of Health Research and Indian Council of Medical Research grant number: 5/13/92/2020/NCD-III awarded to S.K..

## Notes

https://www.ncbi.nlm.nih.gov/bioproject/PRJNA1242580

### Competing Interest Statement

The authors have declared no competing interest.

