## Supplementary figures for "Transcriptomic landscape of Gallbladder cancer reveals altered pathways related to cell cycle and Aurora kinase"

### Figure S1

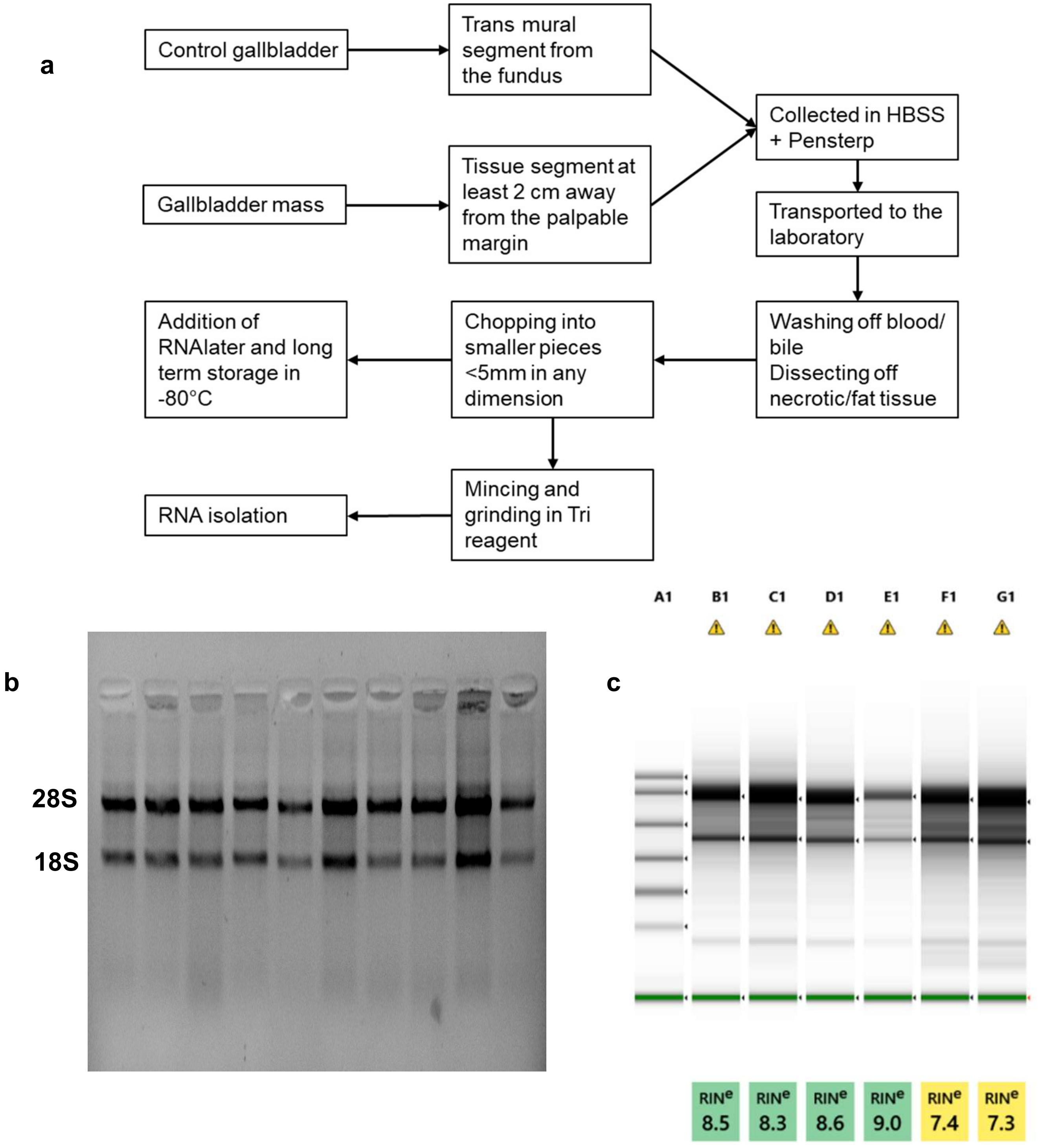


Figure S1: Tissue harvesting and isolation of RNA. a. Workflow for collecting and processing of tissue. b. Representative image of agarose gel electrophoresis and c. RIN scoring by bioanalyzer indicating quality of RNA isolated from tissue samples. HBSS- Hank Basal Salt Solution. Penstrep- Penicillin + Streptomycin.

### Figure S2

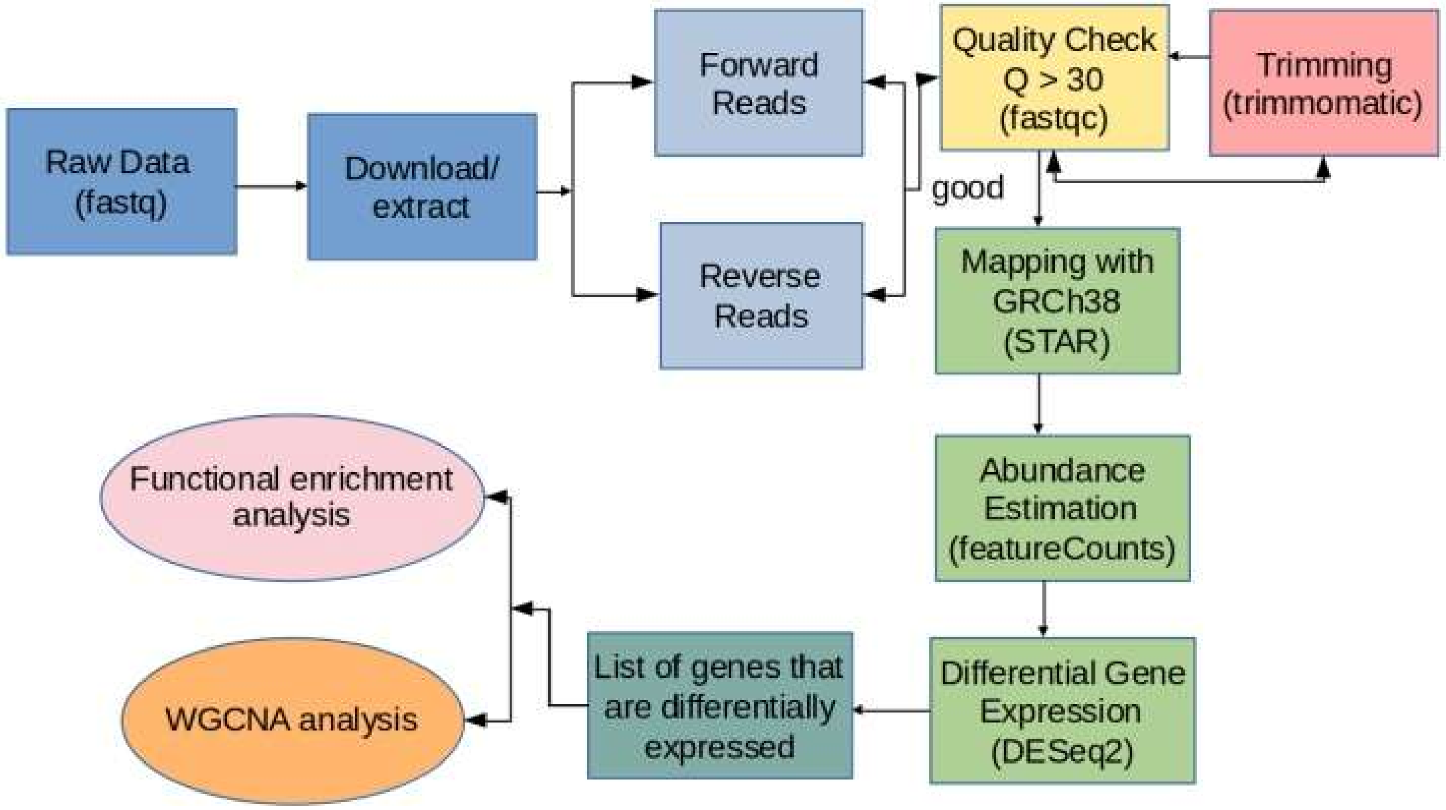


Figure S2: RNA-seq workflow used in the study.

### Figure S3

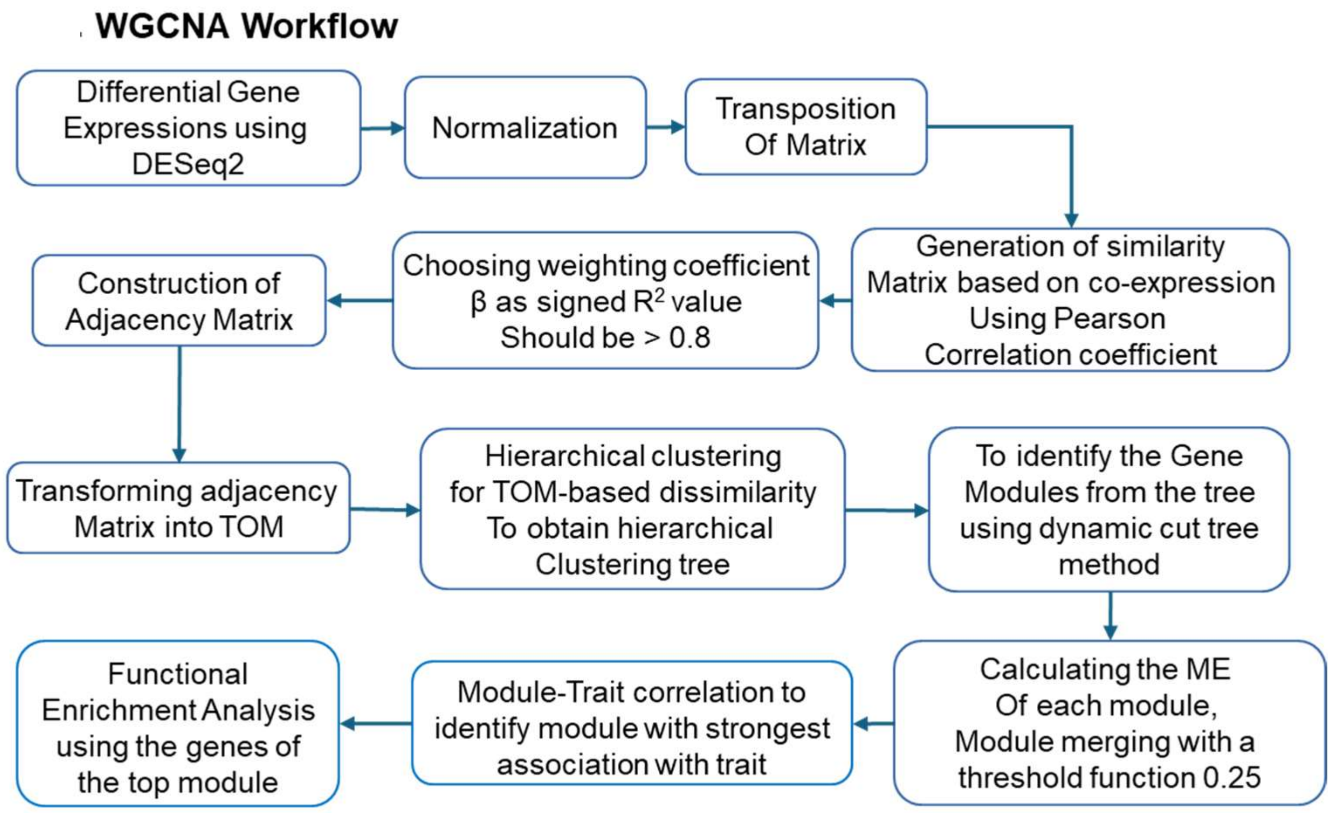


Figure S3: WGCNA workflow used in the study: General outline of the analysis workflow. TOM- Topological overlap Matrix, ME- Module eigengene, GS- Gene significance, MM- Module membership.

### Figure S4

Morphology

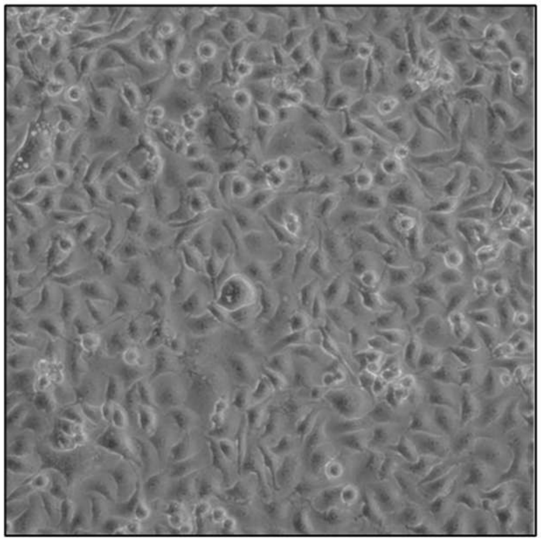

STR profile

| Loci | Database (RCB) | STR profile observed |
|---|---|---|
| Amelogenin | X | X |
| CSF1PO | 12 | 12 |
| D5S818 | 12 | 12 |
| D7S820 | 11,12 | 11,12 |
| D13S317 | 8,10 | 8,10 |
| D16S539 | 9,12 | 9,12 |
| TH01 | 7 | 7 |
| TPOX | 11 | 11 |
| vWA | 16 | 16 |

G415

Figure S4: Morphology and STR profile of the GBC cell line G415 used in the study.

### Figure S5

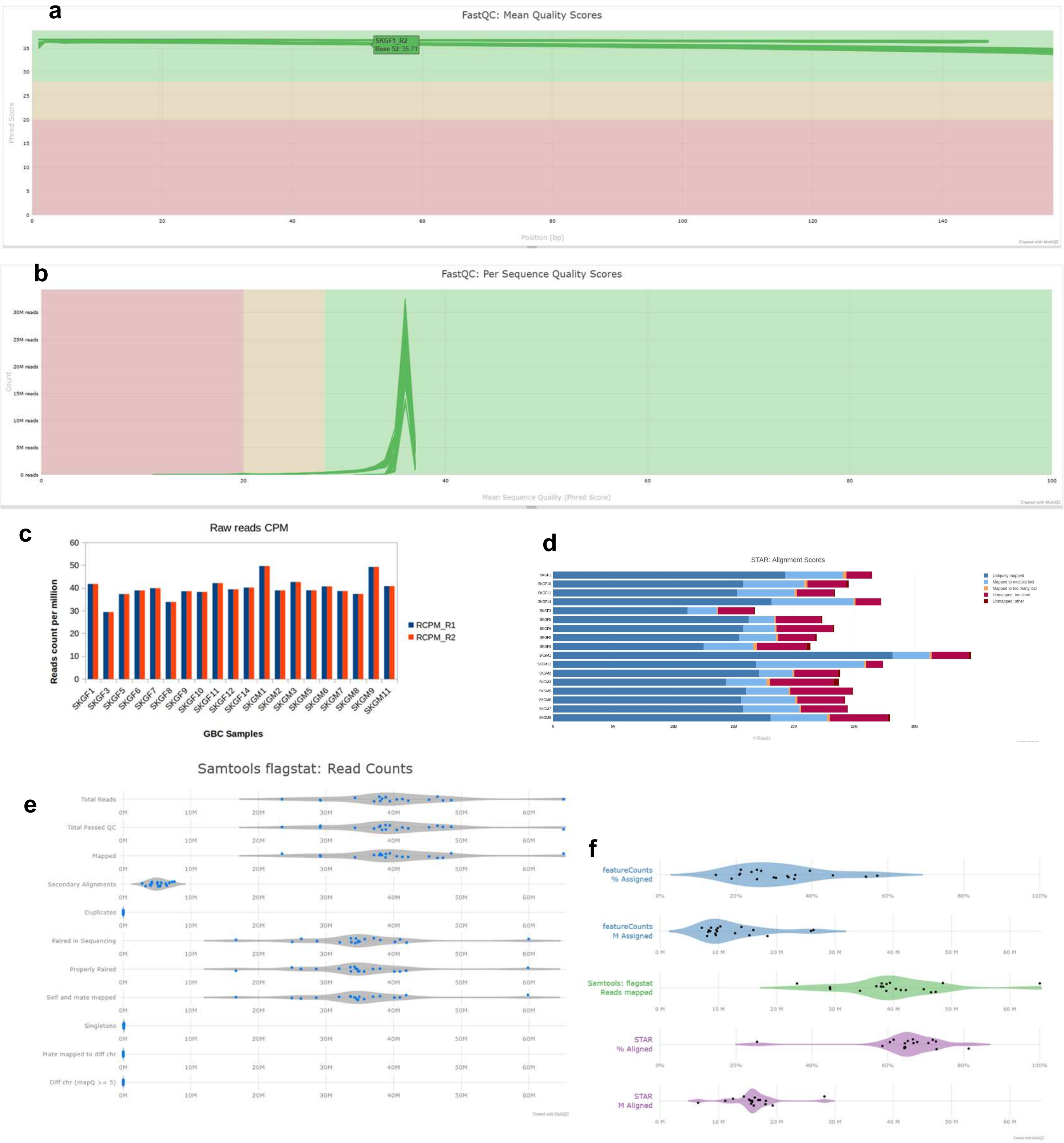


Figure S5: Quality control of the RNAseq data. a. Mean read quality score (Phred score) of different samples. b. Per sequence quality scores of different samples. c. raw read count statistics. d. Alignment statistics as obtained by STAR tool. e. Summary statistics of the raw reads. f. Summary statistics of mapping and alignment.

### Figure S6

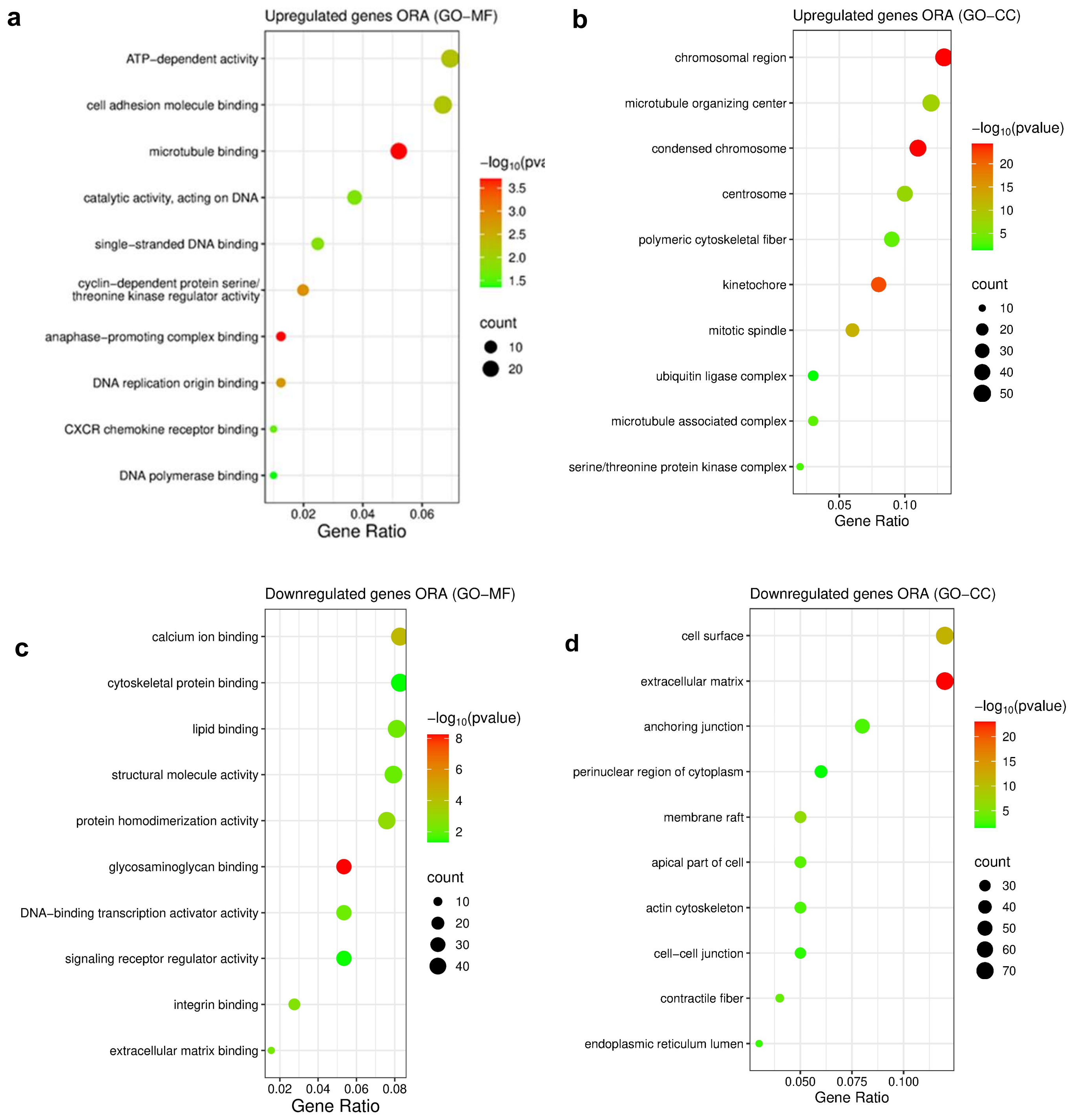


Figure S6: Over Representation analysis of differentially expressed genes using different database. a, b. ORA using upregulated genes respectively with GO-MF and GO-CC database. B & D. ORA using downregulated genes respectively with GO-MF and GO-CC . GO-CC: Gene Ontology- Cellular Components, GO-MF: Gene Ontology- Molecular Function.

### Figure S7

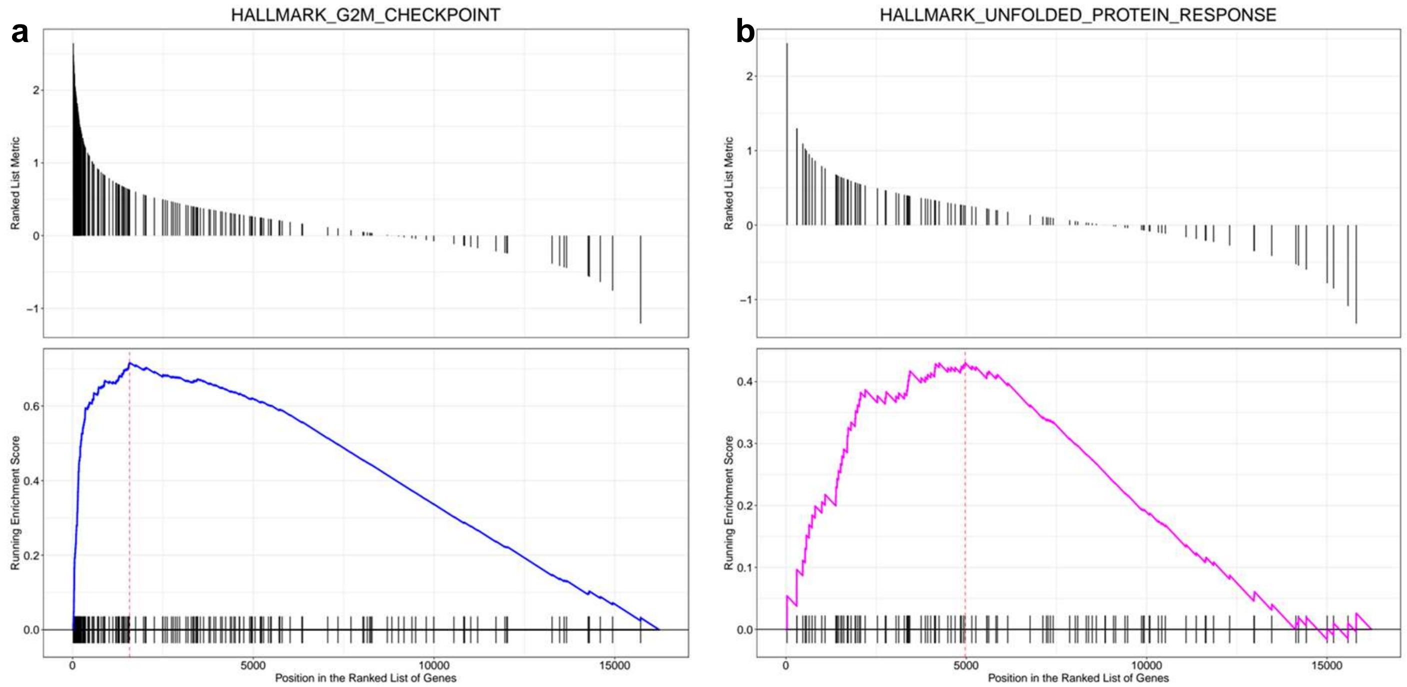


Figure S7: Gene Set Enrichment Analysis of significant pathways. a. Hallmark G2M Checkpoint, b. Hallmark unfolded protein response.

### Figure S8

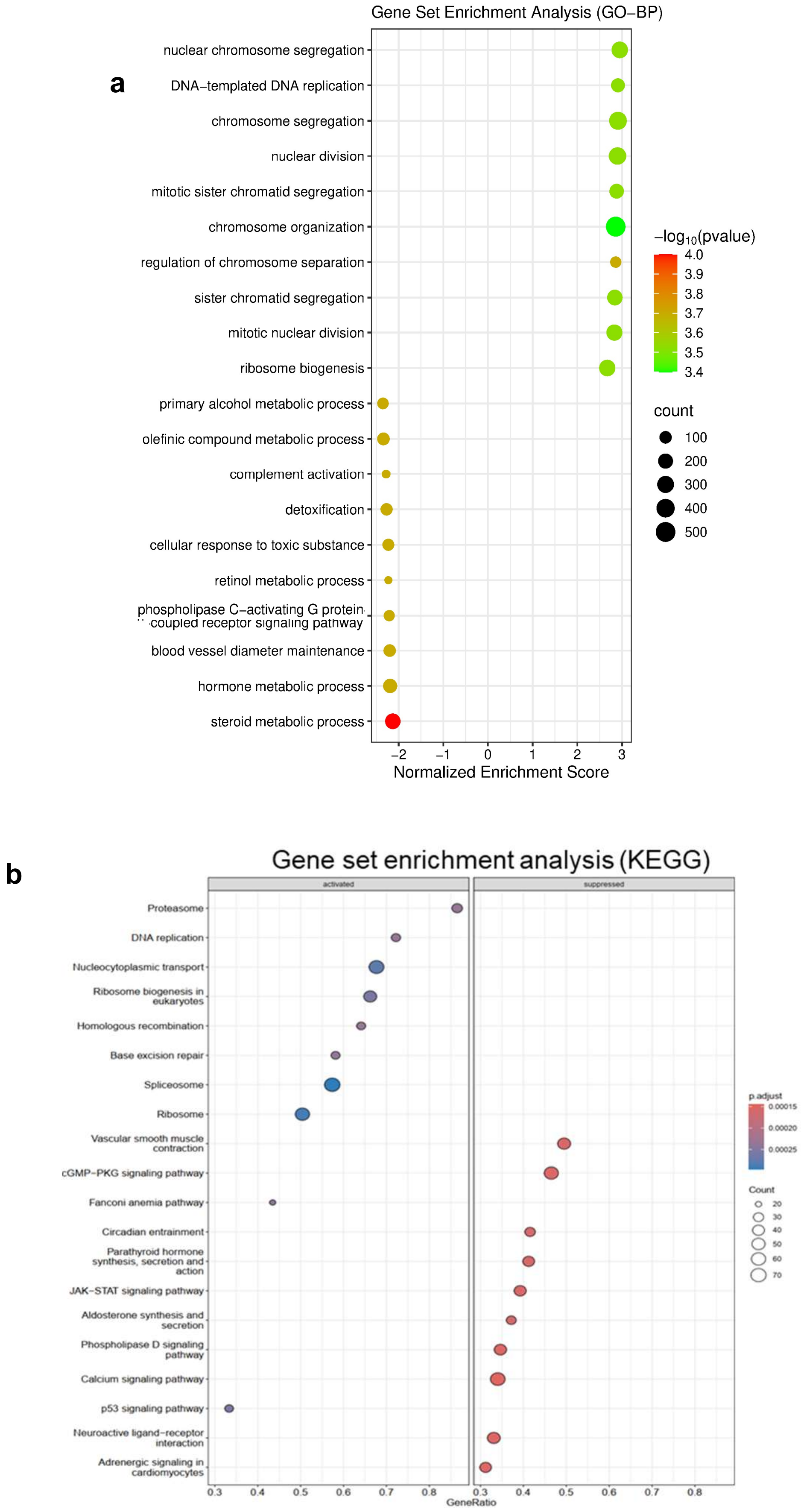


Figure S8: Pathways obtained from Gene Set Enrichment Analysis of differentially expressed genes using different database. a. GSEA with GO-BP database. b. GSEA with KEGG database. GO-BP: Gene Ontology- Biological Processes. KEGG- Kyoto Encyclopedia of Genes and Genomes.

### Figure S9

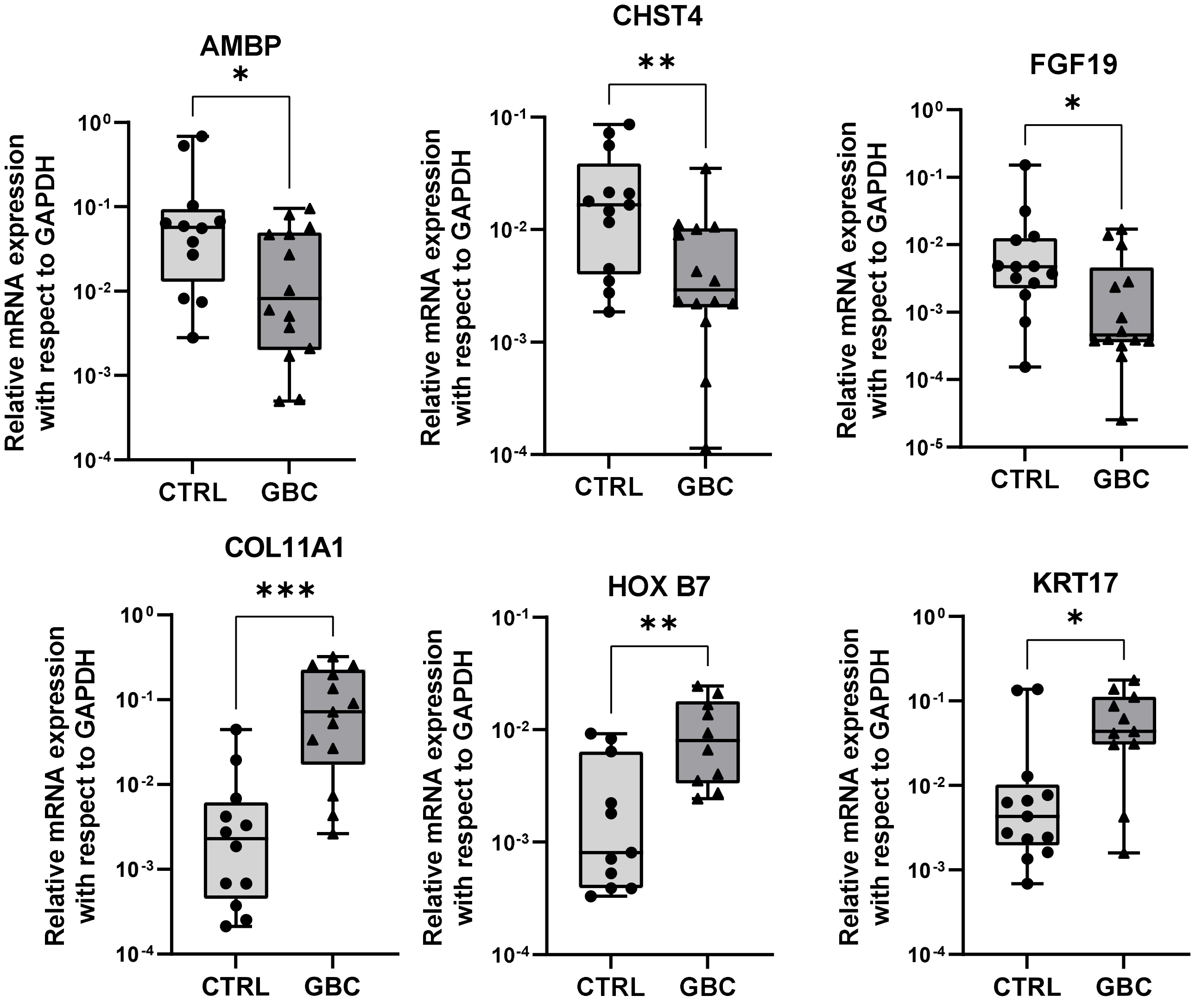


Figure S9: Box plot showing relative mRNA expression of genes (by qPCR).

### Figure S10

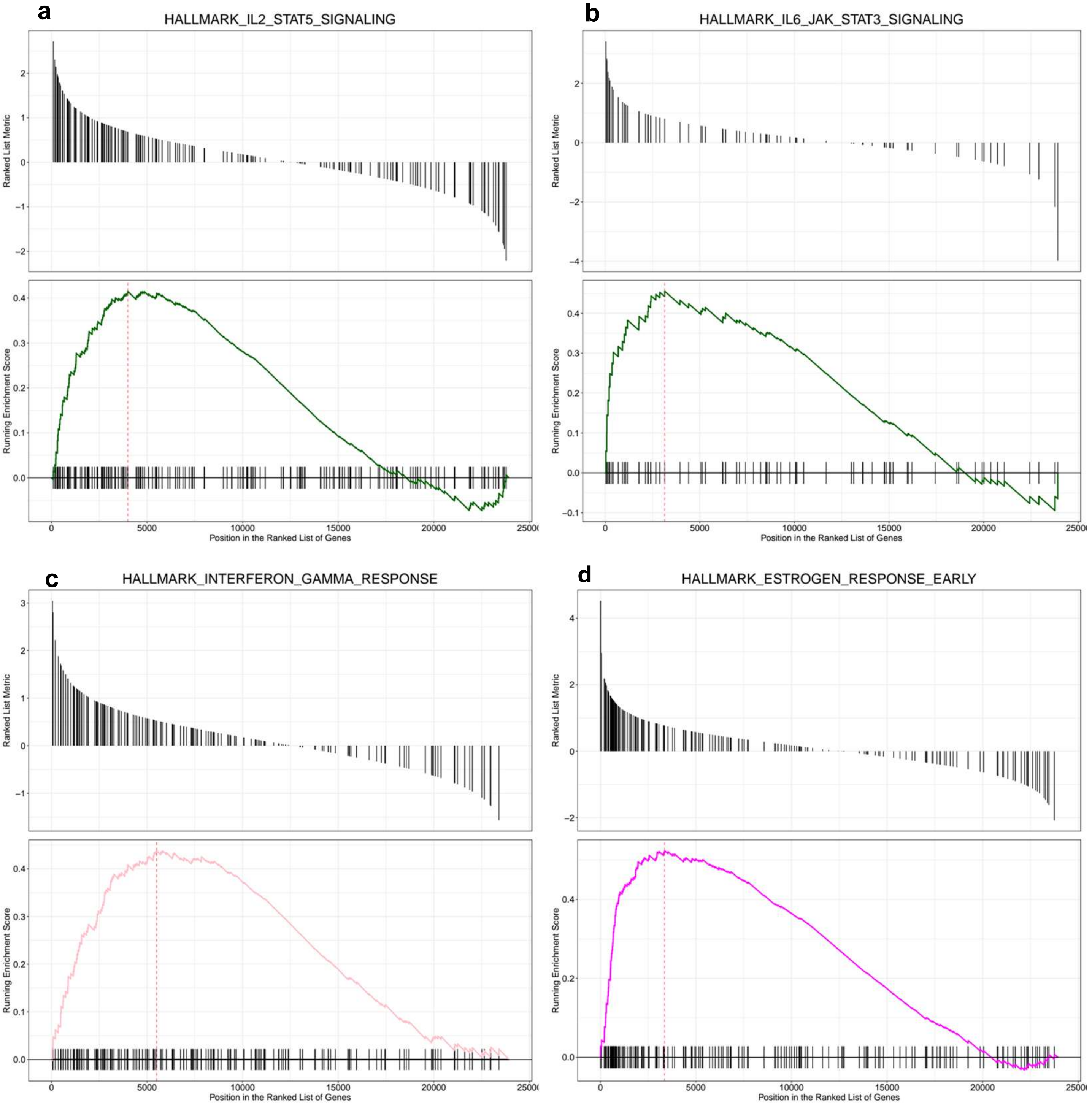


Figure S10: Gene Set Enrichment Analysis of significant pathways in male GBC. a. Halmark IL2-STAT5 signaling, b. Halmark IL6-JAK-STAT3 signaling, c. Hallmark Interferon gamma response. d. Halmark Estrogen response (Early).

### Figure S11

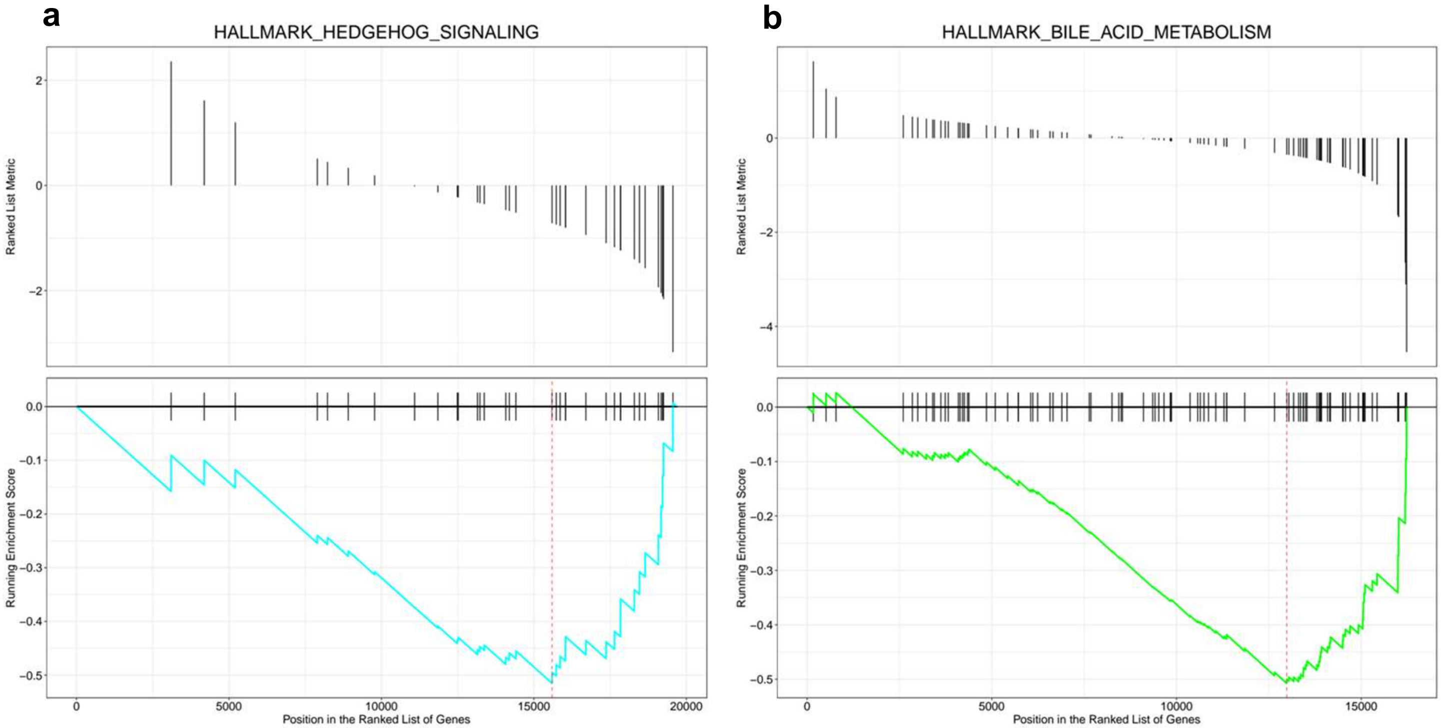


Figure S11: Gene Set Enrichment Analysis of significant pathways in female GBC. a. Halmark Hedgehog signaling. b. Halmark Bile acid metabolism.

### Figure S12

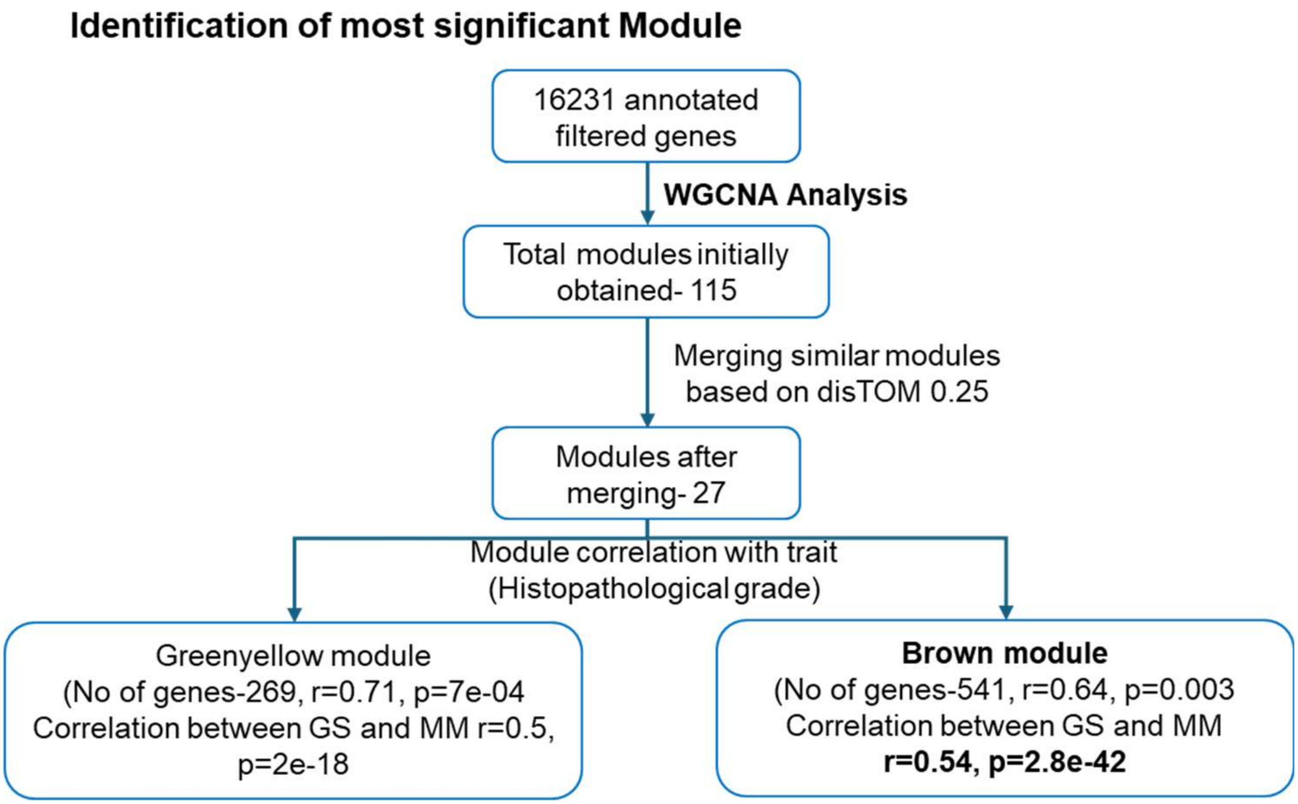


Figure S12: Selection of Brown module by WGCNA for further analysis. ME- Module eigengene, GS- Gene significance, MM- Module membership.

### Figure S13

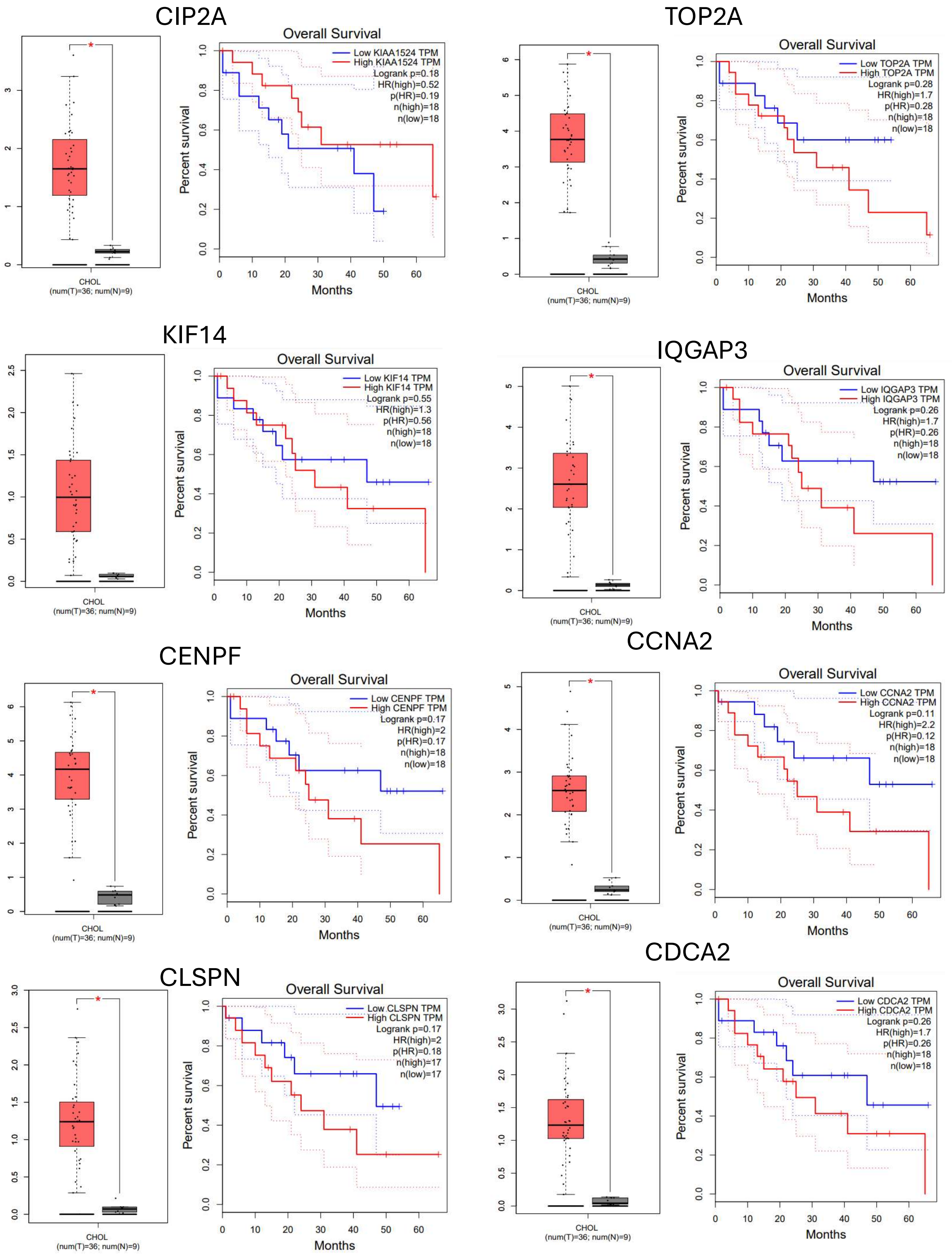


Figure S13: Expression data and overall survival plots of hub genes using TCGA database. Red boxes represent GBC while grey boxes indicate control.

### Figure S14

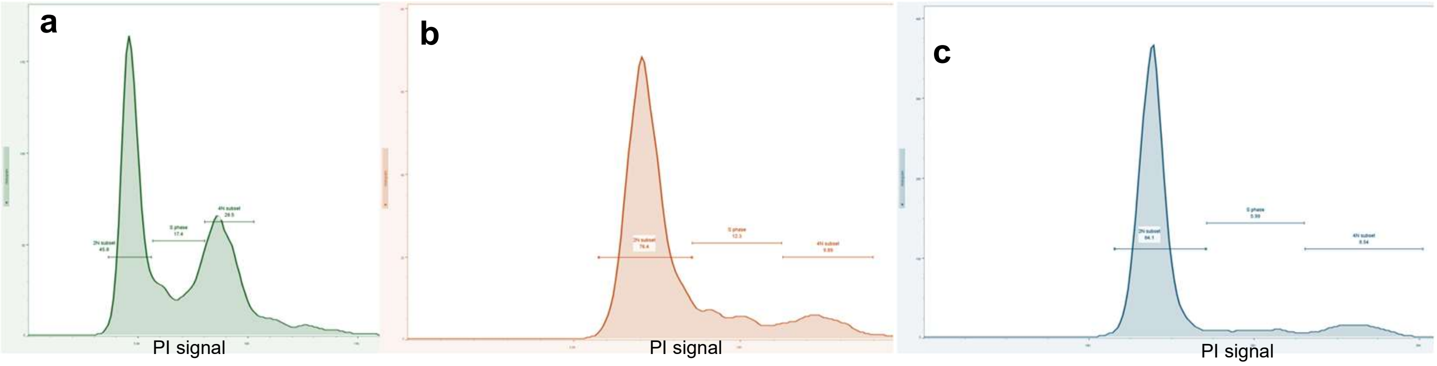


Control

TC-S7010
100nM

Hesperadin
100nM

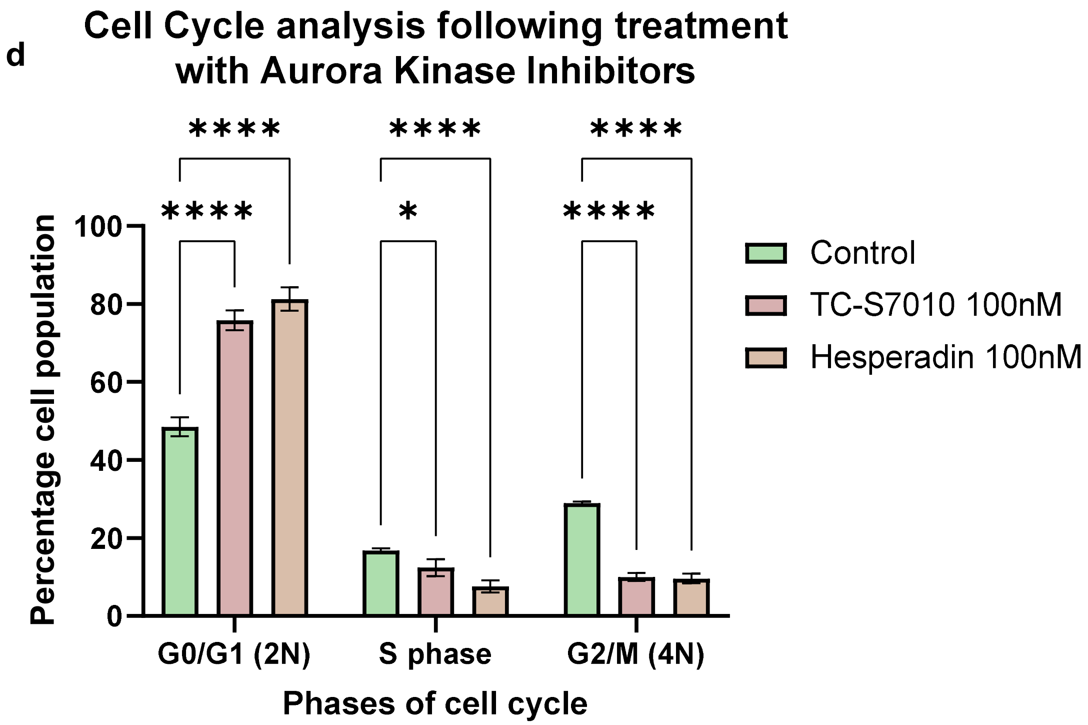


Figure S14: Cell Cycle analysis performed in G415 cell line by propidium iodide staining following treatment with Aurora Kinase Inhibitors. Representative histogram of PI fluorescence of control group (a), AURKA inhibitor TC-S7010 treated cells (b), AURKB inhibitor Hesperadin hydrochloride treated cells (c). Both the treatment was at the dosage of 100nM for 24 hours. **d** Comparison of percentage population of different fractions of cell cycle after treatment with inhibitors (n=3 biological replicates). 2 way ANOVA test was performed followed Dunnet's post hoc test. *p value <0.05, **** p value <0.0001.
