## Supplementary tables for "Transcriptomic landscape of Gallbladder cancer reveals altered pathways related to cell cycle and Aurora kinase"

**Table S1: List of Primers used in the study for qPCR**

| Sl no. | Gene | Forward primer (5->3) | Reverse Primer (5->3) | Product size (base pair) |
|---|---|---|---|---|
| 1 | AMBP | ATGGCTGACCGAGGTGAATG | TCAGTTACCAGTTGCCCACC | 125bp |
| 2 | FGF19 | TTGCTGGAGATCAAGGCAGTC | TCCTCCGAGTACTGAAGCAGC | 119bp |
| 3 | CHST4 | CCTCCCTCAACCTGCATATCG | TCACAATGCGACTGTCAATCA | 105bp |
| 4 | HOXB7 | CGAGCCGAGTTCCTTCAACA | GTTTGCGGTCAGTTCCTGAG | 176bp |
| 5 | Col11A1 | GGCTGAAAGTGTAACAGAGG | TCATCAACGATGTTTGCCTC | 75bp |
| 6 | KRT17 | CCAGATCCAGGGGCTGATTG | TTGTACTGAGTCAGGTGGGC | 177bp |
| 7 | TPX2 | AGGGCCTTTCTGGTTCTCTAGT | GGATTTGCCTTATGCACCAGT | 205bp |
| 8 | CENPF | CGTCCCCGAGAGCAAGTTTA | GTAGGCAGCCCTTCTTTCCA | 98bp |
| 9 | ANLN | GGTGTGGTAAGTCCAGAGAGTT | CACCAGATTCAGCTCGAGGG | 127bp |
| 10 | TOP2A | TGTCACCATTGCAGCCTGT | TGTCTGGGCGGAGCAAAATA | 138bp |
| 11 | PRC1 | CGCCATGAGGAGAAGTGAGG | TCTCTCCTTCTTCCTTGATATGCT | 161bp |
| 12 | KIF23 | TGCCATGAAGTCAGCGAGAG | GCGCACCCTACAGTATACCC | 103bp |
| 13 | GAPDH | ACGGATTTGGTCGTATTGGG | CGCTCCTGGAAGATGGTGAT | 214bp |
| 14 | E-Cadherin | AAAGGCCCATTTCCTAAAAACCT | TGCGTTCTCTATCCAGAGGCT | 172bp |
| 15 | N-Cadherin | CCGGTTTCATTTGAGGGCAC | TTGAGGGCATTGGGATCGTC | 210bp |
| 16 | Vimentin | AGAACTTTGCCGTTGAAGCTG | AGAAATCCTGCTCTCCTCGC | 191bp |
| 17 | MMP2 | AGGATGGCAAGTACGGCTTC | CTTCTTGTCGCGGTCGTAGT | 185bp |
| 18 | MMP9 | GAGCTGACTCGACGGTGATG | AACTGTATCCTTGGTCCGGG | 189bp |
| 19 | TIMP1 | CATTGCTGGAAAACTGCAGGA | GCAGTTTGCAGGGGATGGAT | 167bp |
| 20 | TIMP2 | GGCTGCGAGTGCAAGATCAC | TCGAGAAACTCCTGCTTGGG | 197bp |
| 21 | Ki67 | GCCTGTACGGCTAAAACATGGAG | ACGTGCTGGCTCCTGTTCAC | 133bp |
| 22 | PCNA | GGTTACTGAGGGCGAGAAGC | GACCGGCTGAGACTTGCGTA | 98bp |

| Table S2: List of top 200 differentially expressed genes in GBC compared to control | | | | | |
|---|---|---|---|---|---|
| Upregulated | | | Downregulated | | |
| Sl no | Gene | log2FoldChange | Sl no | Gene | log2FoldChange |
| 1 | COL11A1 | 4.74 | 1 | FGF19 | -7.27 |
| 2 | KRT17 | 3.88 | 2 | AMBP | -5.09 |
| 3 | TNNT1 | 3.79 | 3 | DES | -4.72 |
| 4 | MMP12 | 3.30 | 4 | GC | -4.71 |
| 5 | CHI3L1 | 3.05 | 5 | PI16 | -4.64 |
| 6 | CLDN18 | 3.02 | 6 | SCARA5 | -4.54 |
| 7 | MATN3 | 3.00 | 7 | PAH | -4.39 |
| 8 | CDH3 | 2.94 | 8 | VTN | -4.32 |
| 9 | REG1A | 2.93 | 9 | CHST4 | -4.28 |
| 10 | COL10A1 | 2.92 | 10 | ADH1B | -4.05 |
| 11 | SERPINB5 | 2.89 | 11 | ASGR2 | -3.97 |
| 12 | CXCL10 | 2.86 | 12 | CES1 | -3.96 |
| 13 | LOC101060400 | 2.78 | 13 | AGT | -3.95 |
| 14 | ESM1 | 2.78 | 14 | APCS | -3.93 |
| 15 | NOS2 | 2.77 | 15 | SERPINA5 | -3.84 |
| 16 | CXCL9 | 2.77 | 16 | CRP | -3.84 |
| 17 | TOP2A | 2.75 | 17 | FCGBP | -3.82 |
| 18 | HOXB7 | 2.73 | 18 | IGHA2 | -3.67 |
| 19 | CXCL5 | 2.64 | 19 | MUC5B | -3.64 |
| 20 | HIST1H2BM | 2.62 | 20 | TFF2 | -3.60 |
| 21 | ANLN | 2.61 | 21 | CLEC3B | -3.41 |
| 22 | HIST1H2AH | 2.60 | 22 | RDH12 | -3.38 |
| 23 | POSTN | 2.60 | 23 | ACTG2 | -3.33 |
| 24 | CDC20 | 2.58 | 24 | IGSF10 | -3.33 |
| 25 | HIST1H2AJ | 2.56 | 25 | CLU | -3.26 |
| 26 | IGLV3-21 | 2.55 | 26 | SLC30A2 | -3.22 |
| 27 | HIST1H2AL | 2.52 | 27 | LGALS2 | -3.18 |
| 28 | HIST1H3C | 2.48 | 28 | HABP2 | -3.14 |
| 29 | AURKA | 2.47 | 29 | TTR | -3.12 |
| 30 | HIST1H3B | 2.46 | 30 | UGT2B15 | -3.09 |
| 31 | HOXA13 | 2.46 | 31 | PLP1 | -3.08 |
| 32 | IGHV1-69D | 2.46 | 32 | AKR1B10 | -3.06 |
| 33 | HLA-DRB1 | 2.46 | 33 | SLC15A1 | -3.04 |
| 34 | CENPF | 2.44 | 34 | CA4 | -3.04 |
| 35 | MYBL2 | 2.41 | 35 | KRT222 | -2.95 |
| 36 | KIF26B | 2.41 | 36 | DUOX2 | -2.93 |
| 37 | CCNB2 | 2.40 | 37 | CCKAR | -2.92 |
| 38 | TPX2 | 2.37 | 38 | ALDOB | -2.91 |
| 39 | PLAU | 2.37 | 39 | C16orf89 | -2.84 |
| 40 | FOXM1 | 2.37 | 40 | DIRAS2 | -2.84 |

| | | | | | |
|---|---|---|---|---|---|
| 41 | SULF1 | 2.35 | 41 | GREM2 | -2.82 |
| 42 | ECT2 | 2.35 | 42 | PLIN1 | -2.76 |
| 43 | CDKN3 | 2.33 | 43 | CH25H | -2.76 |
| 44 | UBE2C | 2.33 | 44 | ANXA13 | -2.74 |
| 45 | BIRC5 | 2.31 | 45 | SYNPO2 | -2.72 |
| 46 | NUSAP1 | 2.30 | 46 | TMEM132C | -2.72 |
| 47 | SAPCD2 | 2.28 | 47 | HSPB6 | -2.71 |
| 48 | C5orf46 | 2.27 | 48 | AKR1C4 | -2.70 |
| 49 | FIRRE | 2.26 | 49 | GLP2R | -2.68 |
| 50 | MKI67 | 2.26 | 50 | SERPINA4 | -2.68 |
| 51 | HIST1H2BH | 2.24 | 51 | ALDH1A2 | -2.68 |
| 52 | INHBA | 2.23 | 52 | OGN | -2.67 |
| 53 | HIST1H3F | 2.21 | 53 | ANPEP | -2.67 |
| 54 | BUB1 | 2.21 | 54 | SYNM | -2.66 |
| 55 | CCNE1 | 2.20 | 55 | B3GALT2 | -2.66 |
| 56 | HIST1H3G | 2.19 | 56 | FABP4 | -2.65 |
| 57 | IGKV1-39 | 2.19 | 57 | C6 | -2.64 |
| 58 | LRRC15 | 2.18 | 58 | ADAM33 | -2.61 |
| 59 | PPP1R1B | 2.17 | 59 | PTGDS | -2.59 |
| 60 | HIST1H2BL | 2.16 | 60 | FBLN5 | -2.58 |
| 61 | CEP55 | 2.16 | 61 | CAVIN2 | -2.58 |
| 62 | TYMS | 2.16 | 62 | RBP4 | -2.58 |
| 63 | HIST1H1B | 2.15 | 63 | ABCC2 | -2.58 |
| 64 | NUF2 | 2.14 | 64 | CNN1 | -2.57 |
| 65 | ASPM | 2.14 | 65 | LYVE1 | -2.57 |
| 66 | MELK | 2.13 | 66 | MYOC | -2.55 |
| 67 | BUB1B | 2.13 | 67 | SRPX | -2.55 |
| 68 | HIST1H4I | 2.13 | 68 | COLEC11 | -2.53 |
| 69 | CDK1 | 2.12 | 69 | VSTM2L | -2.53 |
| 70 | ARNTL2 | 2.12 | 70 | C7 | -2.52 |
| 71 | HMMR | 2.12 | 71 | GPX3 | -2.52 |
| 72 | LAMC2 | 2.11 | 72 | CPB2 | -2.51 |
| 73 | LOC105370045 | 2.11 | 73 | CHRDL2 | -2.50 |
| 74 | MMP9 | 2.11 | 74 | CD36 | -2.50 |
| 75 | TRIM29 | 2.11 | 75 | SLC2A4 | -2.50 |
| 76 | DLGAP5 | 2.10 | 76 | AOX1 | -2.49 |
| 77 | LOC105376938 | 2.09 | 77 | HBB | -2.48 |
| 78 | IGLV3-1 | 2.08 | 78 | MAMDC2 | -2.45 |
| 79 | KIF4A | 2.07 | 79 | FGG | -2.43 |
| 80 | MYEOV | 2.07 | 80 | CADM3 | -2.42 |
| 81 | IGLV3-19 | 2.07 | 81 | IGHA1 | -2.41 |
| 82 | LAMB3 | 2.05 | 82 | CNTFR | -2.40 |
| 83 | HIST1H2AM | 2.05 | 83 | HBA2 | -2.40 |
| 84 | HJURP | 2.05 | 84 | MMRN1 | -2.39 |

| | | | | | |
|---|---|---|---|---|---|
| 85 | PLAC8 | 2.05 | 85 | DPT | -2.38 |
| 86 | AFAP1-AS1 | 2.05 | 86 | NGFR | -2.38 |
| 87 | PRC1 | 2.03 | 87 | LMOD1 | -2.37 |
| 88 | HIST1H2AG | 2.02 | 88 | CHRM2 | -2.36 |
| 89 | CDCA8 | 2.02 | 89 | ADAMTS8 | -2.36 |
| 90 | FN1 | 2.02 | 90 | SLC3A1 | -2.35 |
| 91 | S100P | 2.00 | 91 | VIT | -2.34 |
| 92 | GINS2 | 2.00 | 92 | SLC9A3 | -2.34 |
| 93 | ABHD17C | 2.00 | 93 | PODN | -2.32 |
| 94 | MCM10 | 1.99 | 94 | ECM1 | -2.32 |
| 95 | IDO1 | 1.99 | 95 | APOD | -2.32 |
| 96 | TRIP13 | 1.99 | 96 | MUC6 | -2.32 |
| 97 | IL4I1 | 1.98 | 97 | ADGRD1 | -2.30 |
| 98 | HOXA10 | 1.97 | 98 | JCHAIN | -2.30 |
| 99 | HMGB3 | 1.96 | 99 | TM4SF4 | -2.30 |
| 100 | COL8A1 | 1.96 | 100 | GFRA1 | -2.30 |

| Table S3: List of top 200 differentially expressed genes in male GBC compared to control | | | | | |
|---|---|---|---|---|---|
| Upregulated | | | Downregulated | | |
| Sl no | Gene | log2FoldChange | Sl no | Gene | log2FoldChange |
| 1 | COMP | 4.87 | 1 | UGT2B28 | -6.65 |
| 2 | ANXA8L1 | 4.46 | 2 | FABP4 | -5.88 |
| 3 | FNDC1 | 4.30 | 3 | FGF19 | -5.70 |
| 4 | COL8A1 | 4.26 | 4 | PLIN1 | -5.36 |
| 5 | REG3A | 3.81 | 5 | XIST | -5.21 |
| 6 | SERPINE1 | 3.75 | 6 | LOC105375977 | -4.92 |
| 7 | CILP2 | 3.73 | 7 | APCS | -4.43 |
| 8 | VSIG1 | 3.65 | 8 | SFRP5 | -4.24 |
| 9 | MUC2 | 3.61 | 9 | PLIN4 | -4.09 |
| 10 | CCL20 | 3.40 | 10 | CD36 | -3.98 |
| 11 | ANKRD1 | 3.39 | 11 | ADIPOQ | -3.86 |
| 12 | POTEF | 3.32 | 12 | TUSC5 | -3.67 |
| 13 | IGLV3-19 | 3.24 | 13 | UGT2B17 | -3.65 |
| 14 | CLCF1 | 3.20 | 14 | ADAMTS16 | -3.53 |
| 15 | NPTX1 | 3.20 | 15 | CIDEA | -3.40 |
| 16 | IGFBP3 | 3.19 | 16 | EBF2 | -3.37 |
| 17 | ASPHD2 | 3.17 | 17 | RBP7 | -3.31 |
| 18 | BARX2 | 3.09 | 18 | CIDEC | -3.31 |
| 19 | HLA-DRB1 | 3.04 | 19 | ABCA9 | -3.16 |
| 20 | IGKV3-15 | 3.03 | 20 | CLEC4G | -3.11 |
| 21 | CDH2 | 3.01 | 21 | LPL | -3.09 |
| 22 | SPDEF | 2.97 | 22 | CMA1 | -3.08 |
| 23 | SERPINB2 | 2.96 | 23 | PRKAR2B | -3.00 |
| 24 | SFN | 2.96 | 24 | CHST4 | -2.99 |
| 25 | CYP2S1 | 2.94 | 25 | DCLK1 | -2.94 |
| 26 | IGFBP1 | 2.93 | 26 | LRRC2 | -2.93 |
| 27 | IGKV3-20 | 2.92 | 27 | PMP2 | -2.92 |
| 28 | COL8A2 | 2.91 | 28 | THRSP | -2.91 |
| 29 | STC1 | 2.87 | 29 | MYOC | -2.91 |
| 30 | IGKV1-39 | 2.85 | 30 | IGLV2-18 | -2.87 |
| 31 | CXCL3 | 2.85 | 31 | KRT222 | -2.81 |
| 32 | KLHL25 | 2.84 | 32 | CD209 | -2.79 |
| 33 | HMGCS2 | 2.81 | 33 | ITM2A | -2.76 |
| 34 | MFSD2A | 2.80 | 34 | RERGL | -2.68 |
| 35 | IGKC | 2.77 | 35 | AQP7 | -2.65 |
| 36 | CREB3L1 | 2.76 | 36 | RNVU1-18 | -2.62 |
| 37 | ANXA8 | 2.75 | 37 | CYP1A1 | -2.60 |
| 38 | PTPRR | 2.74 | 38 | PTGFR | -2.57 |
| 39 | IGKV1-5 | 2.72 | 39 | LRRC17 | -2.56 |
| 40 | SEMA7A | 2.72 | 40 | CILP | -2.50 |
| 41 | LIF | 2.72 | 41 | SELENOP | -2.49 |
| 42 | FBXO32 | 2.69 | 42 | ABCA10 | -2.48 |
| 43 | FN1 | 2.67 | 43 | LINC02232 | -2.48 |

| | | | | | |
|---|---|---|---|---|---|
| 44 | FXYD3 | 2.66 | 44 | CYP2C8 | -2.47 |
| 45 | GJB2 | 2.65 | 45 | F13A1 | -2.45 |
| 46 | FOSL1 | 2.61 | 46 | MNDA | -2.43 |
| 47 | PGF | 2.60 | 47 | RNU4ATAC | -2.42 |
| 48 | LINC00668 | 2.59 | 48 | CTSG | -2.38 |
| 49 | TMEM158 | 2.59 | 49 | LOC105370205 | -2.32 |
| 50 | RPL10P9 | 2.58 | 50 | LOC107985481 | -2.29 |
| 51 | LAMP5 | 2.55 | 51 | GHR | -2.28 |
| 52 | IGLV3-10 | 2.54 | 52 | LIPE | -2.26 |
| 53 | PRSS21 | 2.51 | 53 | SNCG | -2.25 |
| 54 | CACNG4 | 2.49 | 54 | GPAM | -2.23 |
| 55 | BHLHA15 | 2.48 | 55 | PKHD1L1 | -2.23 |
| 56 | CNNM1 | 2.48 | 56 | XPNPEP2 | -2.22 |
| 57 | LOC107984316 | 2.47 | 57 | LOC102723809 | -2.14 |
| 58 | PKP2 | 2.47 | 58 | LMO3 | -2.13 |
| 59 | THBS2 | 2.47 | 59 | PCOLCE2 | -2.12 |
| 60 | IGLV3-25 | 2.46 | 60 | FLRT2 | -2.12 |
| 61 | SPHK1 | 2.45 | 61 | LRRK2 | -2.09 |
| 62 | SCARF2 | 2.44 | 62 | PRRG3 | -2.08 |
| 63 | GPR3 | 2.43 | 63 | GGTA1P | -2.08 |
| 64 | PROM1 | 2.42 | 64 | CXCL12 | -2.07 |
| 65 | MUC13 | 2.41 | 65 | NRN1 | -2.05 |
| 66 | MDM2 | 2.41 | 66 | LOC105377321 | -2.05 |
| 67 | EPS8L1 | 2.40 | 67 | TLR8 | -1.98 |
| 68 | TACSTD2 | 2.40 | 68 | LINC01697 | -1.98 |
| 69 | PLAC8 | 2.39 | 69 | PLAC9 | -1.96 |
| 70 | LOC100420423 | 2.39 | 70 | SCN9A | -1.95 |
| 71 | RUNDC3A | 2.39 | 71 | HERC2P3 | -1.95 |
| 72 | CGREF1 | 2.38 | 72 | SLC17A4 | -1.94 |
| 73 | FOXD1 | 2.38 | 73 | GYPC | -1.94 |
| 74 | IGLV2-23 | 2.38 | 74 | COL25A1 | -1.93 |
| 75 | CHST2 | 2.38 | 75 | TLR3 | -1.93 |
| 76 | WISP1 | 2.36 | 76 | PTPRD | -1.93 |
| 77 | PYCR1 | 2.34 | 77 | CXXC4 | -1.92 |
| 78 | IGLL5 | 2.34 | 78 | LOC107984428 | -1.90 |
| 79 | PLEKHB1 | 2.34 | 79 | FXYD1 | -1.87 |
| 80 | SLC17A9 | 2.34 | 80 | KLHL4 | -1.85 |
| 81 | PDZK1IP1 | 2.33 | 81 | P2RY13 | -1.84 |
| 82 | PDCD2L | 2.30 | 82 | FAM13C | -1.83 |
| 83 | PMEPA1 | 2.30 | 83 | F8 | -1.83 |
| 84 | CTSE | 2.29 | 84 | C17orf58 | -1.81 |
| 85 | MOXD1 | 2.29 | 85 | LOC105370569 | -1.78 |
| 86 | DYRK2 | 2.28 | 86 | SEMA3C | -1.77 |
| 87 | TSPAN1 | 2.28 | 87 | FMO2 | -1.76 |
| 88 | MS4A8 | 2.28 | 88 | WISP2 | -1.75 |
| 89 | TNFRSF11B | 2.27 | 89 | WDFY3-AS2 | -1.73 |
| 90 | HIST1H2AL | 2.27 | 90 | ACKR4 | -1.71 |
| 91 | OLR1 | 2.26 | 91 | EMCN | -1.71 |

| | | | | | |
|---|---|---|---|---|---|
| 92 | FAM83D | 2.26 | 92 | EPHA7 | -1.71 |
| 93 | IGLV3-1 | 2.26 | 93 | ARHGAP20 | -1.69 |
| 94 | GPAT2 | 2.25 | 94 | TIMP4 | -1.69 |
| 95 | MICAL2 | 2.24 | 95 | RADIL | -1.69 |
| 96 | MAP7D2 | 2.23 | 96 | MMP16 | -1.69 |
| 97 | RGPD2 | 2.21 | 97 | CLIC2 | -1.68 |
| 98 | SLC16A10 | 2.20 | 98 | PDE1C | -1.67 |
| 99 | IL17RB | 2.19 | 99 | IGDCC4 | -1.67 |
| 100 | ARNT2 | 2.19 | 100 | GUSBP4 | -1.65 |

| Table S4: List of top 200 differentially expressed genes in Female GBC compared to control | | | | | |
|---|---|---|---|---|---|
| Upregulated | | | Downregulated | | |
| Sl no | Gene | log2FoldChange | Sl no | Gene | log2FoldChange |
| 1 | LOC105372647 | 10.70 | 1 | AMBP | -7.43 |
| 2 | SOX11 | 9.62 | 2 | DES | -6.88 |
| 3 | WDR87 | 9.60 | 3 | PI16 | -6.85 |
| 4 | FAM133A | 9.58 | 4 | VTN | -6.71 |
| 5 | CDX2 | 9.57 | 5 | GC | -6.68 |
| 6 | MAGEA12 | 9.29 | 6 | MUC5B | -6.54 |
| 7 | SCEL | 9.28 | 7 | LGALS2 | -6.47 |
| 8 | HOXC10 | 9.15 | 8 | AGT | -5.76 |
| 9 | PASD1 | 9.12 | 9 | PAH | -5.68 |
| 10 | NMU | 9.08 | 10 | AKR1B10 | -5.41 |
| 11 | LINC01705 | 9.03 | 11 | DUOX2 | -5.18 |
| 12 | EPYC | 9.01 | 12 | GREM2 | -5.15 |
| 13 | HOXA13 | 8.95 | 13 | ALDOB | -5.13 |
| 14 | LOC101060400 | 8.93 | 14 | CRP | -5.13 |
| 15 | LOC105371028 | 8.92 | 15 | SLC9A3 | -5.12 |
| 16 | CACNA1S | 8.92 | 16 | AKR7A3 | -5.11 |
| 17 | DSG3 | 8.83 | 17 | SERPINA5 | -5.09 |
| 18 | SLC6A14 | 8.82 | 18 | UGT2B15 | -5.08 |
| 19 | LOC100287912 | 8.73 | 19 | ACTG2 | -5.06 |
| 20 | LOC105374798 | 8.72 | 20 | SLC13A1 | -5.00 |
| 21 | LOC102723623 | 8.69 | 21 | HABP2 | -4.99 |
| 22 | LOC105376361 | 8.69 | 22 | CES1 | -4.94 |
| 23 | LOC105375026 | 8.69 | 23 | TTR | -4.94 |
| 24 | ATP1B4 | 8.64 | 24 | ANXA13 | -4.93 |
| 25 | GOLGA6C | 8.64 | 25 | CLU | -4.92 |
| 26 | LOC105370890 | 8.63 | 26 | ASGR2 | -4.89 |
| 27 | LOC105376980 | 8.62 | 27 | SLC30A2 | -4.87 |
| 28 | PCDH11X | 8.62 | 28 | CHIT1 | -4.86 |
| 29 | LHFPL3-AS2 | 8.62 | 29 | SLC15A1 | -4.83 |
| 30 | EN2 | 8.56 | 30 | FCGBP | -4.68 |
| 31 | HOXD13 | 8.53 | 31 | CST4 | -4.66 |
| 32 | GOLGA6L1 | 8.52 | 32 | GSTA2 | -4.59 |
| 33 | PAX2 | 8.50 | 33 | C4A | -4.56 |
| 34 | TRIML2 | 8.50 | 34 | CA4 | -4.54 |
| 35 | KRT17 | 8.49 | 35 | SCARA5 | -4.54 |
| 36 | LOC112268221 | 8.48 | 36 | ADRA2A | -4.50 |
| 37 | LOC105374062 | 8.48 | 37 | RDH12 | -4.47 |
| 38 | LOC105372577 | 8.46 | 38 | CNN1 | -4.39 |
| 39 | SPANXB1 | 8.44 | 39 | ACE2 | -4.37 |
| 40 | LINC01605 | 8.43 | 40 | PLA2G2A | -4.36 |
| 41 | LOC105372639 | 8.43 | 41 | HSPB6 | -4.32 |
| 42 | LOC105371077 | 8.41 | 42 | GP2 | -4.32 |

| | | | | | |
|---|---|---|---|---|---|
| 43 | EPHA8 | 8.39 | 43 | VSTM2L | -4.30 |
| 44 | SPHKAP | 8.38 | 44 | GSTA1 | -4.28 |
| 45 | TAF1L | 8.38 | 45 | PP7080 | -4.28 |
| 46 | DMBX1 | 8.37 | 46 | SYT8 | -4.27 |
| 47 | PAX7 | 8.37 | 47 | MADCAM1 | -4.26 |
| 48 | SNAR-C3 | 8.37 | 48 | LCN2 | -4.24 |
| 49 | LOC105371466 | 8.37 | 49 | ALDH1A2 | -4.22 |
| 50 | LRRC74B | 8.37 | 50 | CACNA1H | -4.15 |
| 51 | CCDC190 | 8.36 | 51 | HLA-DRB4 | -4.14 |
| 52 | LPA | 8.34 | 52 | FAM107A | -4.12 |
| 53 | LINC01206 | 8.33 | 53 | CLDN2 | -4.11 |
| 54 | ITIH6 | 8.31 | 54 | IGHA2 | -4.10 |
| 55 | GAGE10 | 8.30 | 55 | ONECUT1 | -4.09 |
| 56 | HOTTIP | 8.30 | 56 | SOX17 | -4.07 |
| 57 | LOC105374270 | 8.30 | 57 | ABCC2 | -4.06 |
| 58 | FAM71E2 | 8.29 | 58 | DUOXA2 | -4.05 |
| 59 | LOC105376412 | 8.28 | 59 | COLEC11 | -4.02 |
| 60 | BAGE2 | 8.28 | 60 | ADH1B | -4.01 |
| 61 | LOC101060022 | 8.27 | 61 | ANPEP | -3.98 |
| 62 | LOC105370455 | 8.27 | 62 | RBP4 | -3.98 |
| 63 | DNAJB8 | 8.27 | 63 | SULT1C4 | -3.96 |
| 64 | C2orf54 | 8.25 | 64 | TMEM86B | -3.92 |
| 65 | LOC105377473 | 8.25 | 65 | HPN | -3.92 |
| 66 | LYPD4 | 8.25 | 66 | FBLN5 | -3.91 |
| 67 | CASC8 | 8.24 | 67 | NGFR | -3.87 |
| 68 | LOC105374357 | 8.22 | 68 | PTX3 | -3.86 |
| 69 | LINC01839 | 8.20 | 69 | SYNM | -3.86 |
| 70 | FCRL4 | 8.19 | 70 | TFR2 | -3.83 |
| 71 | LOC105376620 | 8.18 | 71 | PDZK1 | -3.83 |
| 72 | IRS4 | 8.18 | 72 | FOSB | -3.83 |
| 73 | HOXB13 | 8.17 | 73 | AOX1 | -3.83 |
| 74 | LOC105376214 | 8.17 | 74 | ECM1 | -3.81 |
| 75 | OPRK1 | 8.17 | 75 | UCA1 | -3.79 |
| 76 | LOC105372026 | 8.16 | 76 | KCNJ16 | -3.78 |
| 77 | LOC105372549 | 8.16 | 77 | LOC101928277 | -3.76 |
| 78 | LOC112268183 | 8.15 | 78 | PODN | -3.71 |
| 79 | LINC01789 | 8.14 | 79 | SYNPO2 | -3.70 |
| 80 | GPR87 | 8.14 | 80 | LOC105369230 | -3.69 |
| 81 | ARGFX | 8.14 | 81 | C8G | -3.68 |
| 82 | SIX3 | 8.13 | 82 | MUC20 | -3.67 |
| 83 | LINC02286 | 8.11 | 83 | CYP4F11 | -3.66 |
| 84 | PRDM9 | 8.11 | 84 | AKR1C4 | -3.66 |
| 85 | PRSS47 | 8.11 | 85 | CLEC3B | -3.65 |
| 86 | PRDM7 | 8.10 | 86 | WFDC1 | -3.63 |
| 87 | HOXA11 | 8.10 | 87 | CASR | -3.62 |
| 88 | HOXC6 | 8.10 | 88 | TM4SF4 | -3.61 |

| | | | | | |
|---|---|---|---|---|---|
| 89 | IL36RN | 8.10 | 89 | MUC3A | -3.59 |
| 90 | SLCO1B3 | 8.10 | 90 | LMOD1 | -3.59 |
| 91 | LOC107984019 | 8.09 | 91 | KCNK5 | -3.56 |
| 92 | NCAN | 8.05 | 92 | GPX3 | -3.56 |
| 93 | FABP7 | 8.05 | 93 | LINC00261 | -3.56 |
| 94 | ESPNP | 8.05 | 94 | PCK1 | -3.55 |
| 95 | LOC105372252 | 8.04 | 95 | KLK11 | -3.55 |
| 96 | KIRREL2 | 8.03 | 96 | PIGR | -3.54 |
| 97 | TEX13D | 8.02 | 97 | SLC4A4 | -3.52 |
| 98 | C17orf77 | 8.02 | 98 | GABRB3 | -3.50 |
| 99 | FAR2P1 | 8.01 | 99 | ZBTB16 | -3.50 |
| 100 | LOC105378901 | 8.01 | 100 | SERPINA4 | -3.49 |

| Table S5: Filtered genelist of Brown module | | | | | |
|---|---|---|---|---|---|
| **Sl no** | **Gene** | **Intramodular Connectivity** | **Module Membership** | **Gene Significance with respect to histopathological grade** | **q.Weighted** |
| 1 | TPX2 | 89.41 | 0.97 | 0.70 | 2.06512E-06 |
| 2 | CENPF | 89.05 | 0.95 | 0.63 | 4.44402E-05 |
| 3 | KIF14 | 88.20 | 0.96 | 0.53 | 0.00065634 |
| 4 | KIF4A | 88.09 | 0.97 | 0.60 | 3.66168E-05 |
| 5 | TOP2A | 87.24 | 0.95 | 0.68 | 5.70629E-06 |
| 6 | IQGAP3 | 87.17 | 0.97 | 0.61 | 5.66237E-05 |
| 7 | CLSPN | 86.14 | 0.97 | 0.55 | 0.000324351 |
| 8 | ANLN | 84.52 | 0.96 | 0.73 | 1.00551E-06 |
| 9 | CENPE | 83.77 | 0.95 | 0.54 | 0.000564134 |
| 10 | BUB1B | 83.67 | 0.96 | 0.70 | 4.45494E-06 |
| 11 | CIP2A | 82.85 | 0.96 | 0.60 | 0.000143054 |
| 12 | KIF23 | 82.75 | 0.96 | 0.67 | 9.70972E-06 |
| 13 | CENPK | 82.32 | 0.95 | 0.64 | 5.93586E-05 |
| 14 | MELK | 81.96 | 0.96 | 0.71 | 2.85036E-06 |
| 15 | BUB1 | 80.68 | 0.95 | 0.68 | 3.35347E-06 |
| 16 | CCNA2 | 80.36 | 0.96 | 0.65 | 3.34346E-05 |
| 17 | CDCA2 | 80.10 | 0.96 | 0.58 | 0.000152649 |
| 18 | DLGAP5 | 78.86 | 0.94 | 0.60 | 7.92341E-05 |
| 19 | PRC1 | 78.75 | 0.94 | 0.72 | 2.5268E-06 |
| 20 | ATAD2 | 78.48 | 0.95 | 0.63 | 4.60368E-05 |
| 21 | NUSAP1 | 77.39 | 0.94 | 0.68 | 1.78832E-05 |
| 22 | CDK1 | 76.97 | 0.94 | 0.65 | 1.9699E-05 |
| 23 | NDC80 | 76.73 | 0.94 | 0.65 | 1.65137E-05 |
| 24 | HMMR | 75.36 | 0.93 | 0.66 | 1.53966E-05 |
| 25 | GINS1 | 73.77 | 0.94 | 0.59 | 7.60458E-05 |
| 26 | KIF20A | 73.58 | 0.95 | 0.66 | 4.99718E-06 |
| 27 | CKAP2L | 73.35 | 0.93 | 0.60 | 0.000572492 |
| 28 | HJURP | 72.71 | 0.94 | 0.67 | 6.38002E-06 |
| 29 | KNL1 | 72.28 | 0.92 | 0.63 | 0.000214049 |
| 30 | CCNB2 | 72.07 | 0.93 | 0.72 | 2.46936E-06 |
| 31 | BRCA1 | 71.43 | 0.93 | 0.55 | 0.000693886 |
| 32 | MCM10 | 71.31 | 0.94 | 0.65 | 1.53506E-05 |
| 33 | BIRC5 | 71.22 | 0.94 | 0.67 | 7.73277E-06 |
| 34 | DTL | 69.00 | 0.93 | 0.63 | 0.000101134 |
| 35 | NCAPG | 68.72 | 0.92 | 0.53 | 0.000337832 |
| 36 | KIF18B | 68.51 | 0.94 | 0.55 | 0.000513909 |
| 37 | NUF2 | 68.16 | 0.91 | 0.56 | 0.00037325 |
| 38 | TTK | 66.83 | 0.91 | 0.54 | 0.000430871 |
| 39 | ERCC6L | 66.16 | 0.91 | 0.58 | 0.000582435 |
| 40 | MAD2L1 | 65.05 | 0.91 | 0.59 | 0.000118123 |
| 41 | STIL | 64.88 | 0.92 | 0.54 | 0.00098206 |
| 42 | SMC4 | 64.38 | 0.91 | 0.68 | 5.48344E-05 |
| 43 | KIF2C | 64.11 | 0.92 | 0.61 | 2.20342E-05 |
| 44 | KIF11 | 63.90 | 0.90 | 0.61 | 7.24916E-05 |
| 45 | CDC45 | 63.89 | 0.93 | 0.59 | 0.000279628 |
| 46 | CDCA5 | 63.59 | 0.93 | 0.66 | 5.59662E-06 |
| 47 | CCNB1 | 62.80 | 0.91 | 0.65 | 5.14958E-06 |
| 48 | ZWILCH | 62.76 | 0.89 | 0.65 | 0.000101202 |
| 49 | FANCI | 62.70 | 0.91 | 0.62 | 7.4789E-06 |
| 50 | CDC6 | 62.66 | 0.91 | 0.60 | 7.76839E-05 |
| 51 | MKI67 | 62.62 | 0.90 | 0.73 | 2.3183E-06 |
| 52 | PRR11 | 61.58 | 0.92 | 0.68 | 9.09196E-06 |
| 53 | RAD51 | 59.42 | 0.92 | 0.69 | 1.38099E-05 |

| | | | | | |
|---|---|---|---|---|---|
| 54 | GTSE1 | 59.30 | 0.92 | 0.54 | 0.000496528 |
| 55 | RRM2 | 58.97 | 0.91 | 0.61 | 9.42316E-05 |
| 56 | CDCA8 | 58.62 | 0.90 | 0.64 | 4.12299E-05 |
| 57 | UBE2C | 58.26 | 0.90 | 0.66 | 4.57159E-06 |
| 58 | CDCA3 | 57.52 | 0.92 | 0.51 | 0.000634342 |
| 59 | SHCBP1 | 56.79 | 0.91 | 0.68 | 9.45053E-06 |
| 60 | TYMS | 56.78 | 0.91 | 0.66 | 6.08279E-06 |
| 61 | PTTG1 | 56.22 | 0.90 | 0.61 | 9.36924E-05 |
| 62 | CEP55 | 55.47 | 0.90 | 0.66 | 6.14739E-05 |
| 63 | DSCC1 | 54.51 | 0.91 | 0.52 | 0.000621294 |
| 64 | RACGAP1 | 53.93 | 0.91 | 0.51 | 0.000460381 |
| 65 | ZWINT | 53.64 | 0.89 | 0.65 | 9.93971E-06 |
| 66 | MYBL2 | 53.62 | 0.90 | 0.73 | 6.62329E-07 |
| 67 | KIF20B | 53.53 | 0.86 | 0.54 | 0.000417414 |
| 68 | SPC24 | 53.17 | 0.90 | 0.58 | 3.14509E-05 |
| 69 | GINS2 | 53.08 | 0.90 | 0.71 | 1.79176E-05 |
| 70 | ECT2 | 53.01 | 0.89 | 0.72 | 1.32721E-05 |
| 71 | E2F1 | 51.62 | 0.90 | 0.61 | 8.92546E-05 |
| 72 | TRIP13 | 51.59 | 0.88 | 0.67 | 2.2741E-05 |
| 73 | CDKN3 | 51.34 | 0.87 | 0.57 | 0.000686317 |
| 74 | AURKB | 51.15 | 0.90 | 0.61 | 4.76601E-05 |
| 75 | ORC6 | 49.49 | 0.88 | 0.62 | 2.65506E-05 |
| 76 | CKS2 | 48.42 | 0.86 | 0.68 | 4.35213E-06 |
| 77 | MND1 | 47.55 | 0.87 | 0.61 | 0.000355088 |
| 78 | CDC20 | 46.84 | 0.89 | 0.65 | 5.11964E-06 |
| 79 | RFC4 | 44.41 | 0.87 | 0.60 | 4.06878E-05 |
| 80 | PLK1 | 44.39 | 0.87 | 0.66 | 2.27822E-06 |
| 81 | FOXM1 | 43.46 | 0.87 | 0.70 | 1.39231E-06 |
| 82 | SERPINB5 | 42.31 | 0.86 | 0.71 | 0.000287179 |
| 83 | SPAG5 | 42.07 | 0.88 | 0.60 | 0.000499254 |
| 84 | CENPL | 40.73 | 0.84 | 0.52 | 0.000682689 |
| 85 | BLM | 40.51 | 0.84 | 0.50 | 0.000805984 |
| 86 | TEDC2 | 37.07 | 0.86 | 0.60 | 0.000195915 |
| 87 | POC1A | 35.90 | 0.85 | 0.60 | 0.000281627 |
| 88 | CENPW | 33.34 | 0.82 | 0.59 | 0.000320019 |
| 89 | SAPCD2 | 31.18 | 0.83 | 0.62 | 3.61038E-05 |
| 90 | UHRF1 | 30.99 | 0.83 | 0.71 | 1.87973E-05 |
| 91 | TK1 | 30.24 | 0.82 | 0.73 | 1.39991E-05 |
| 92 | FBXO5 | 29.01 | 0.81 | 0.62 | 0.000702208 |
| 93 | TOPBP1 | 28.94 | 0.77 | 0.65 | 0.000419795 |
| 94 | SPDL1 | 28.60 | 0.79 | 0.64 | 0.000618976 |
| 95 | LMNB1 | 28.35 | 0.82 | 0.58 | 0.000511461 |
| 96 | CCNF | 27.83 | 0.82 | 0.56 | 0.000807126 |
| 97 | CHEK2 | 27.15 | 0.83 | 0.56 | 0.000219645 |
| 98 | CDC25A | 22.33 | 0.78 | 0.55 | 0.000884401 |
| 99 | LOC101060400 | 21.75 | 0.77 | 0.73 | 0.000122762 |
| 100 | MASTL | 21.43 | 0.75 | 0.59 | 0.000979849 |
| 101 | NUDCD1 | 21.07 | 0.77 | 0.72 | 1.21885E-05 |
| 102 | ASCC3 | 20.79 | 0.73 | 0.67 | 7.40858E-05 |
| 103 | HASPIN | 20.08 | 0.77 | 0.50 | 0.000355337 |
| 104 | ARNTL2 | 19.37 | 0.77 | 0.73 | 2.27322E-05 |
| 105 | CDCA4 | 17.91 | 0.75 | 0.63 | 0.000301733 |
| 106 | DBF4 | 17.60 | 0.73 | 0.63 | 0.00021547 |
| 107 | MYEOV | 17.43 | 0.76 | 0.57 | 0.000386671 |
| 108 | C11orf80 | 15.61 | 0.74 | 0.53 | 0.000922748 |
| 109 | CCNE1 | 14.76 | 0.72 | 0.65 | 0.000338794 |
| 110 | ERI1 | 6.76 | 0.61 | 0.82 | 1.68957E-05 |
